# Optimizing and assessing multichannel TMS focality

**DOI:** 10.1101/2025.09.19.677136

**Authors:** Ole Numssen, Carla W. Martin, Torge Worbs, Axel Thielscher, Konstantin Weise, Thomas R. Knösche

## Abstract

**Background:** Multichannel transcranial magnetic stimulation (mTMS) enables electronic steering of induced electric fields across multiple cortical targets without physical coil repositioning, addressing key limitations of conventional single-channel TMS (sTMS). However, determining optimal input currents for focal stimulation remains challenging, and different mTMS systems have not been systematically compared under realistic hardware constraints.

**Objective:** To develop an user-centric framework for optimizing and assessing mTMS focality by introducing a generic optimization algorithm, establishing meaningful quality metrics, and comparing mTMS coil arrays with traditional single-channel TMS across cortical targets.

**Methods:** We developed a fast optimization framework incorporating target E-field constraints via parametrization of degenerated hyperellipsoids, explicitly integrating current-rate limits, for example from stimulator electronics and coil heating. Using high-resolution finite-element models of nine individual brains, we compared two mTMS designs (5-channel planar and 6/12-channel spherical systems) with standard sTMS figure-of-eight coils through E-field simulations. Three complementary metrics quantified stimulation quality: Focality, Target2Max, and OverstimulatedArea.

**Results:** Despite using a single optimized placement for all region-of-interest targets, mTMS achieved focality comparable to repositioned single-channel TMS in this *in-silico* study. For superficial targets, single-channel TMS showed slightly better stimulation, but for deeper cortical targets (>25 mm skin-cortex distance), mTMS performed similarly. More mTMS channels improved focality but required stronger current-rate constraints. The planar design performed better for deeper targets, while spherical designs improved with additional channels.

**Conclusion:** mTMS systems demonstrate remarkable performance comparable to standard TMS, enabling efficient multi-target stimulation without repositioning. Our open-source framework provides practical tools for designing and evaluating mTMS systems, supporting goal-directed mTMS development and effective application.

## Introduction

Transcranial Magnetic Stimulation (TMS) is a non-invasive brain stimulation technique that has revolutionized our ability to investigate causal functional relationships in the human cortex. By inducing electric fields (E-fields) in cortical tissue, TMS can directly trigger action potentials, enabling effective perturbation of neural circuits. This capability complements correlative neuroimaging techniques, such as fMRI or EEG, by allowing causal inference in brain function. Conventional single-channel TMS (sTMS), particularly when applied at the cortical activation threshold, has demonstrated high spatial precision in motor and visual regions (Thielscher et al., 2010; Siebner et al., 2022). However, targeting different cortical regions sequentially can only be achieved by physically moving the sTMS coil, which is a time-consuming and error-prone process (Souza et al., 2025a; Numssen et al., 2024). Additionally, the perceived focality of sTMS from motor cortex studies can be treacherous, as motor cortex targets might often be circumscribed regions residing on the gyral crowns and are therefore easy to target differentially. Realizing this precision in more complex cortical areas remains a challenge, as the shape of the induced E-field from sTMS cannot be changed (Numssen et al., 2023).

Recent advancements in TMS hardware (Koponen et al., 2018, Navarro de Lara et al., 2021) have introduced multi-channel or multi-locus TMS (mTMS) as a promising solution to these limitations. Unlike traditional sTMS, mTMS integrates multiple coils (channels) with different shapes in one multi-channel coil array, enabling channels to be driven independently with individual current rates and directions. The resulting total E-field (E_total_) is a superposition of the single-channel E-fields (E_single_) and can be shaped electronically by tuning the single-channel current rates (Fig. 1a; Daneshzand et al., 2025a; Souza et al., 2022). Importantly, this innovation allows for modifying the spatial stimulation profile, that is adjusting peak location, orientation (Souza et al., 2025a), and spread (Daneshzand et al., 2025b), without the need for physical coil repositioning (Nieminen et al., 2022; Nieminen et al., 2019). This capability facilitates electronic targeting of stimulation hotspots, enabling rapid, sequential modulation of multiple cortical areas with millisecond temporal resolution. Such temporal precision is crucial for probing network dynamics, studying causal interactions across distributed brain regions, and advancing time-sensitive neuromodulation therapies (Sinisalo et al., 2024a). While the concept of superimposing multiple E-fields has been proposed for some time (Ruohonen & Ilmoniemi, 1998), mTMS remains in a developmental phase. At present, several mTMS designs are under investigation, including a 5-channel surface coil array (Koponen et al., 2018; Nieminen et al., 2022) utilizing a layered arrangement of planar coils to optimize field distribution, and a design employing modular units with 3 orthogonally oriented coils each (Navarro de Lara et al., 2021; Daneshzand et al., 2025a) (Fig. 2b). While these approaches have been designed with different use-case in mind, such as fine-grained control over the E-field orientation versus modularity and whole-brain coverage, both are based on the concept of superpositioning several single-channel E-fields to yield one, customizable E_total_ field to allow for electronic steering of the cortical stimulation pattern.

**Figure 1.**
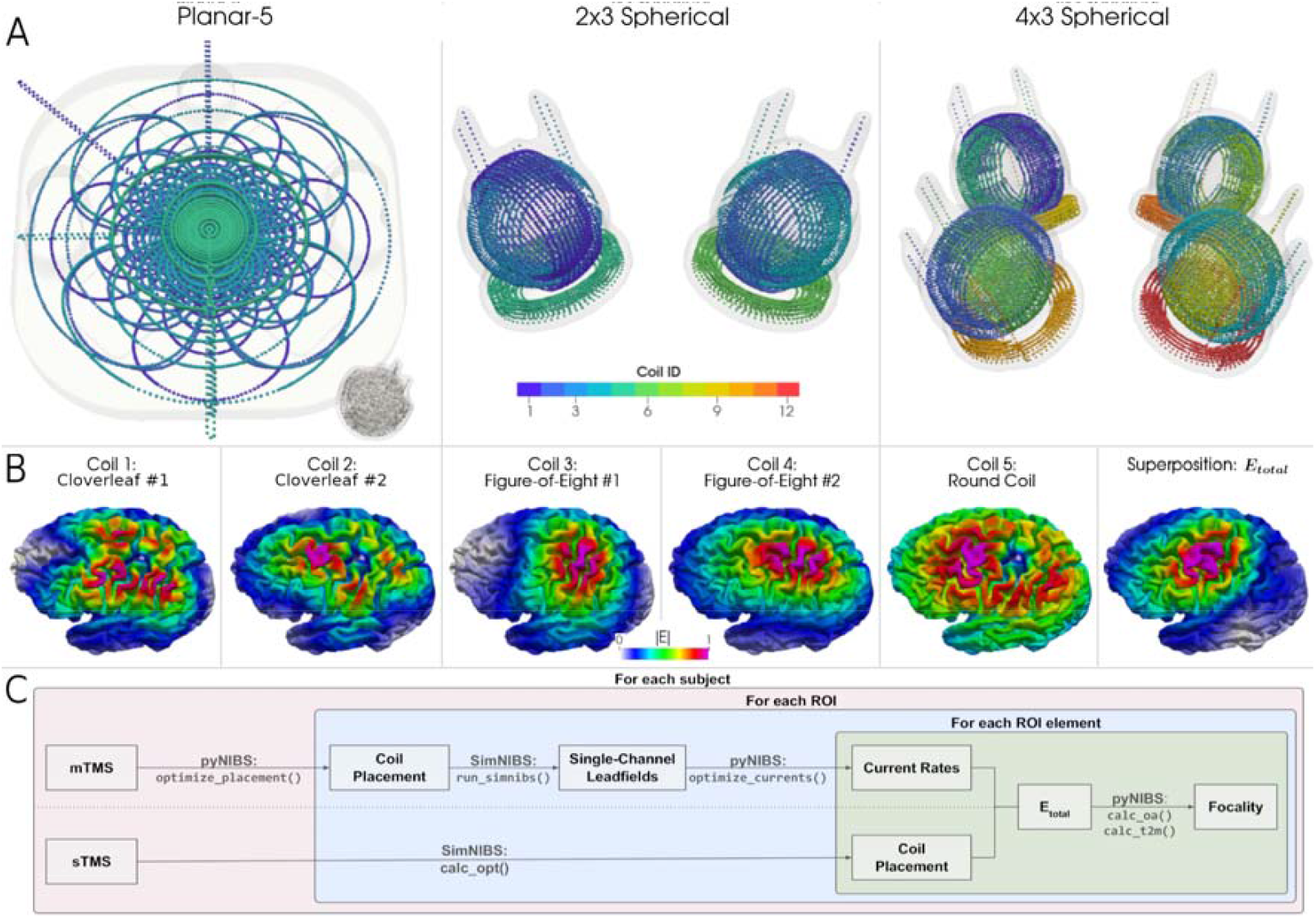
Focality optimization and assessment for single-channel (sTMS) and multi-channel TMS (mTMS). A) We constructed models for three mTMS coil arrays: A planar design consisting of five planar coils (left); and two modular, spherical designs, consisting of two (middle) and four (right) three-channel units, respectively, with one circular coil per x-, y-, and z-axis. Inset in left panel: A three-axis unit for size comparison. B) Single-channel E-fields examples for the 5-chan planar coil. After multiplying these lead fields with the actual current rates *j*, i.e., the stimulator output, their vector-superposition yields the total induced E-field E_total_ (right). C) Focality optimization pipeline. After head-and-brain mesh construction from individual MRI, grey-matter midlayer region of interests (ROIs) were built for the primary motor cortex region for nine healthy individuals across the adult life span (mean±SD age: 53.1 ± 26.80). For mTMS, one coil placement was identified to optimally stimulate the handknob region and single-channel E-fields (‘lead fields’) were computed. Subsequently, for 100 random ROI elements, the optimal current rates, i.e., the stimulator output, were determined for focal stimulation. In contrast, for sTMS, the coil placements were optimized for each of the same 100 ROI elements according to the current sTMS state of the art. Finally, focality was assessed for mTMS and sTMS coils for these targets.

**Figure 2.**
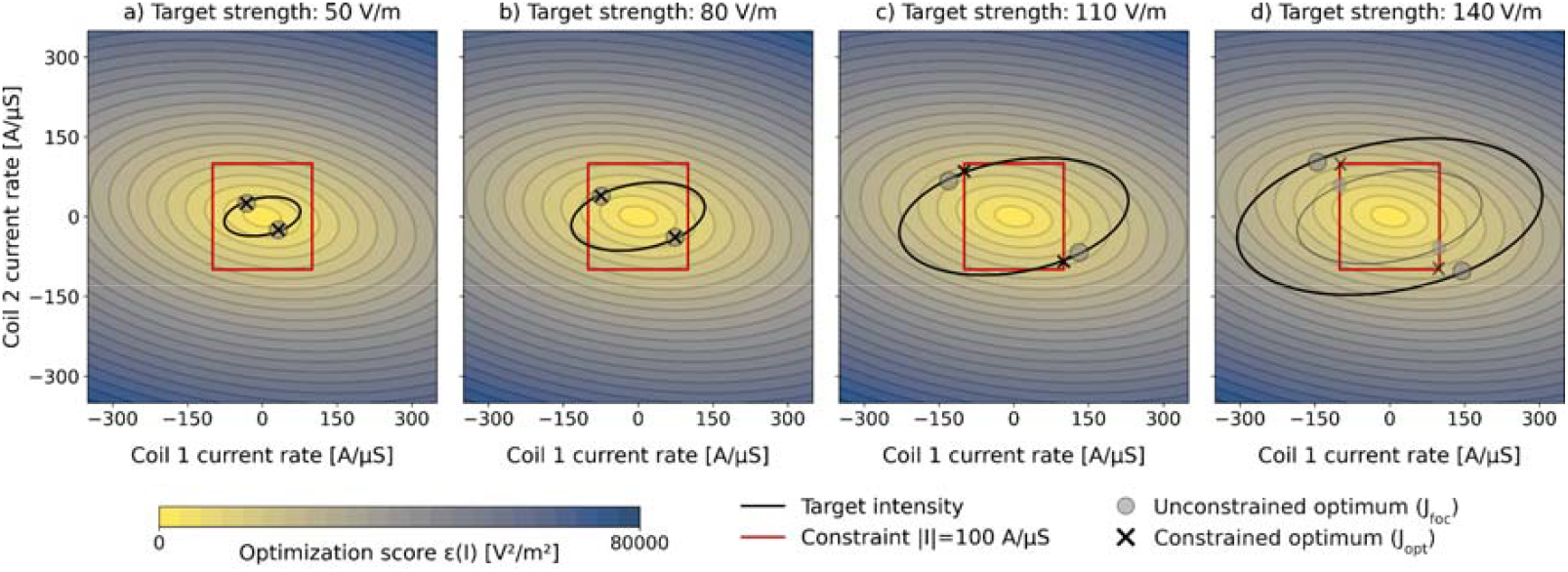
Different optimization cases, illustrated with a hypothetical two-channel coil array. The focality optimization minimizes E-field magnitudes across all cortical locations, while maintaining the target E-field magnitude at the cortical target (minimizing, see Eq. 10). a– b) For low target E-field magnitude or targets close to the mTMS array, optimal solutions (grey circles) lie within the current-rate constraints (red rectangle, *J*_*foc*_). c) At higher target E-field magnitude, these constraints limit the solution space, leading to suboptimal focality (black crosses, *J*_*opt*_). d) Some target E-field magnitudes are not feasible with the current constraints. In these cases, the target E-field magnitude needs to be reduced. Optimal solutions (black crosses) are on some corners of the hypercube, i.e., the absolute values of all channel currents are at the limit.

These promising advancements raise the question of how spatially selective mTMS systems perform in realistic stimulation scenarios, both in comparison to one another and to standard figure□of□eight sTMS coils. This stimulation *focality* is defined here as the ability to deliver a sufficient E-field magnitude at specific cortical target, while suppressing the E-field at off-target regions. Besides maximizing focality, realistic current rate constraints need to be considered, stemming from practical limitations such as capacitor size and heat dissipation.

This study systematically addresses this question by developing and evaluating a framework for optimizing and assessing mTMS focality. Specifically, we (a) introduce a generic and adaptable focality optimization algorithm that can be applied to any mTMS system. Specifically, we (a) introduce a generic and adaptable focality optimization algorithm that can be applied to any mTMS system. It is designed to evenly minimize the off-target stimulation (by minimizing the L2-norm of the E-field vector), while keeping the target stimulation strength constant. Importantly, we reparametrize this optimization goal function to provide a computationally fast and robust solution. (b) For further characterization of the obtained solutions, we develop meaningful stimulation quality metrics for realistic quantification of stimulation precision (*Target2Max*) and off-target spread (*OverstimulatedArea*) for individual head and brain morphology and realistic stimulation strengths. (c) We compare the optimized stimulation patterns of mTMS coil arrays with traditional single-channel sTMS across cortical targets, assessing stimulation efficacy and precision in both superficial and deep cortical areas, and (d) quantify the loss of focality required to yield effective (i.e., sufficiently strong) stimulation strength across cortical regions for a set of mTMS systems.

Throughout this study, we refer to *channel* as an independently drivable wiring (coil), a *coil array* as the physical device which contains all channels in a fixed geometrical arrangement and is placed on the head to generate the electro-magnetic field, and a *stimulator* as the device generating the currents to drive the single channels in the coil array (Fig. 1b).

By providing a framework for focality optimization which is theory-driven, practically sound, easy to use, and adaptable to future mTMS system, this work allows to characterize mTMS systems in a user-centered manner by answering the question “*What is the maximum focality for effective stimulation of a given cortical target?*” With this, we lay the foundation for more refined and controlled stimulation paradigms, fully exploiting currently available as well as future mTMS hardware.

## Methods

### Study Design

This study employed *in silico* experiments to optimize and evaluate the spatial focality of two major multi-channel TMS (mTMS) designs: a 5 channel planar multi-coil design (*5-chan planar:* Fig. 1a, left; Koponen et al., 2018) and a modular 3-axis design with two and four 3-axis spherical units yielding 6 and 12 channels, respectively (*6-chan spherical & 12-chan spherical*; Fig. 1a, center & right; Navarro de Lara et al., 2021). For each mTMS system, we constructed coil models using SimNIBS 4.5 (Worbs et al., 2025), based on publicly available specifications for the spherical systems and on direct communication with the developers for the planar system. To allow comparison with sTMS coils, the MagVenture Cool-B35 and MCF-B65 coils (Drakaki et al., 2022) were selected as representative figure-of-eight coils, commonly used in TMS research and clinical applications. E-field simulations were performed using high-resolution individual head models (SimNIBS 4.5; Thielscher et al., 2015) derived from structural MRI data of nine healthy individuals (mean±SD age: 53.1±26.80) across the lifespan from CamCAN repository (https://cam-can.mrc-cbu.cam.ac.uk/dataset/), based on the CamCAN study (Shafto et al., 2014) and a spherical head model. See SI for details on coil models and head meshes.

### E-field simulation

We determined the mTMS and sTMS placements for stimulating a gyral crown target in the handknob motor region, as a typical TMS target (Ridding et al., 1997; Barker et al., 1985). For the mTMS coil arrays we selected the placement that optimized stimulation focality (see Eq. 10) in the center of the region of interest (ROI) (see SI for details) and used these placements (one per mTMS coil array and subject) throughout the focality analysis. In contrast, for the sTMS coils, we identified one coil placement that maximized the E-field magnitude (|E|) for each cortical target within the ROI (for each sTMS coil and subject), as this is the only parameter that can be varied with sTMS.

E-field simulations were conducted using the SimNIBS toolbox (v4.5; Thielscher et al., 2025) and the pyNIBS Python package (v0.2026.1; Numssen et al., 2021). For each coil array, the lead field matrix *L* was computed, with single-channel E-fields for unit strength current rate (Fig. 1b). The ROI was defined at the cortical gray matter mid-layer to mitigate tissue boundary artifacts for the realistic head models and at the grey matter surface for the spherical head models. The lead field matrices were multiplied with vectors of realistic channel current rates (in A/µs) to derive the final E-field distributions (in V/m) (Jing et al., 2024). Mutual inductance effects, potentially occurring during stimulation in real-word scenarios, were ignored for the purpose of this simulation study.

### Focality Optimization

We aim to increase stimulation focality by maximizing the difference in E-field magnitude between target and off-target regions through the optimization of the current rates in the channels. The whole ROI E-field can be described by concatenating all single-element E-field vectors into a single vector, where is the number of elements (locations) in the ROI. *E* can be computed as the superposition of E-fields, each generated by a unity current rate (e.g., 1 A/µs) in the respective channel, and multiplied with channel-wise current rates. These single-channel E-fields form the columns of the lead field matrix *L* ∈ *R*^*3N_elms* × *N_chans*^ and are scaled by the elements of the current rate vector *J* ∈ *R*^*N_chans*^ :

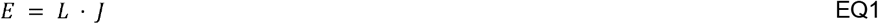

By minimizing the sum of the squares of all E-field magnitudes in the ROI (equivalent to minimizing |*E*|^2^), while keeping the E-field magnitude at the target location |*E*_*target*_| (stimulation strength) fixed to a desired value *e*, we maximize stimulation focality:

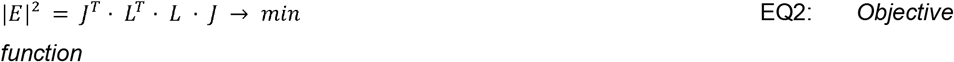

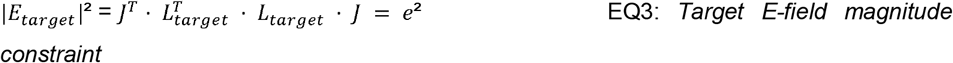

with *L*_*target*_ ∈ *R*^*3*×*N_chans*^ being the lead field for the target location.

Currently, there is no consensus on the exact E-field magnitude required for effective stimulation. Additionally, this stimulation threshold might depend on a set of factors such as the target region, functional context, and brain state (Numssen et al., 2024). A typical E-field magnitude observed at motor threshold intensity is *e*=100 V/m (Weise et al., 2025; Caulfield et al., 2021), which will be used as a standard value in this study. All presented methods are equally valid with any other stimulation threshold.

Note that equal values of the objective function (Eq. 2) form contours, which are hyper-ellipsoids in *R*^*N_chans*^, while Eq. 3 defines a degenerated hyper-ellipsoid (ellipsoid-hypercylinder) surface in the same space, with three finite axes (for the three E-field components in the target). The minima of the objective function on the surface of this degenerated hyper-ellipsoid are the solution of the problem^1^, if we ignore any constraints regarding the maximum possible current rates in the channels.

Such current rate constraints would reflect the limits of the stimulator electronics as well as heat dissipation and mechanical stability in the coil array. They can be interpreted as a hyper-cube (if equal in all channels, otherwise a hyper-cuboid) in *R*^*N_chans*^, restricting the solution to its interior. We can distinguish three cases (see Fig. 2a for a geometrical illustration of the problem for a two-channel coil array): 1) The solutions of Eq. 2 and Eq. 3 are inside the hyper-cube. In this case, the current rate constraints can be disregarded (Fig. 2a-b); 2) The surface of the ellipsoid-hypercylinder is completely outside the hyper-cube, rendering any solution obeying Eq. 3 impossible, that is, the desired E-field magnitude e cannot be reached while keeping all the current rates within the limits. In this case, the best possible solution is the one that maximizes the target E-field magnitude, which corresponds to some corners of the hyper-cube (Fig. 2d); 3) The hyper-cube and the ellipsoid-hypercylinder intersect such that the optimal solutions of Eq. 2 and Eq. 3 are not inside the hyper-cube. In this case, the optimization of Eq. 2 has to be performed only on the parts of the surface of the ellipsoid-hypercylinder that are inside the hyper-cube (Fig. 2c).

### Parameterization

Because the optimization is constrained to the surface of the ellipsoid-hypercylinder defined by Eq. 3, we parameterize that surface. By means of SVD of the lead field matrix at the target, the axes of this degenerated hyperellipsoid can be identified as columns of defined as follows:

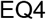

*∑*_*target*_ is a diagonal matrix, where the first three diagonal elements contain the inverses of the squared half-axes of the ellipsoid, while the remaining ones are zero, accounting for the cylindrical part.

We can now insert Eq. 4 into Eq. 3 and obtain an axis-parallel ellipsoid-hypercylinder in the rotated coordinates *X* defined by *U*_*target*_:

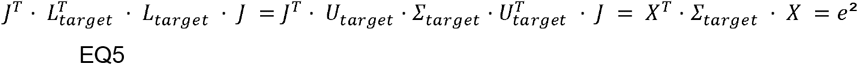

This surface can now be parameterized using the standard formula for an ellipsoidal surface: *X* = *f* (*ϕ*) with angular parameters 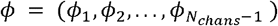 representing locations on the surface of the ellipsoid-hypercylinder:

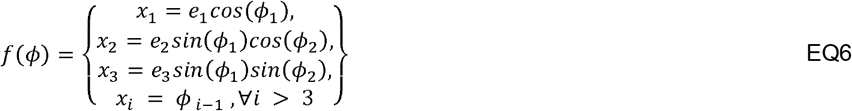

where *e*_1_,*e*_2_,*e*_3_ are the non-zero scaling factors (half-axes lengths) derived from *∑*_*target*_ and 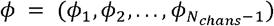 are the angular coordinates of the ellipsoid-hypercylinder.

### Goal function

According to Eq. 5, the parameterization can be linked to the current rates in the channels:

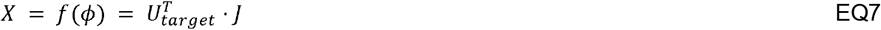

In contrast to Eq. 3, the rank of Eq. 7 is full. Hence, it can be inverted to get an expression for the current rates:

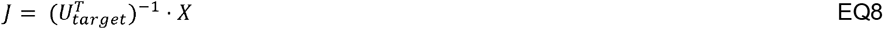

Because *U*_*target*_ is an orthogonal matrix, this equals to:

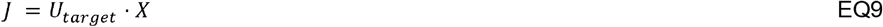

By inserting the parameterization Eq. 7 into Eq. 9 and the result into the objective function Eq. 2, we incorporated the constraint Eq. 3 analytically. This yields an unconstrained nonlinear optimization problem where the minimization parameters are the angular variables *ϕ*_*i*_ :

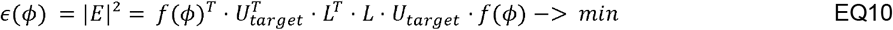

In this formulation, the vector □ parameterizes the surface of the ellipsoid-hypercylinder. The optimization algorithm seeks the solution that minimizes |E| according to Eq. 2, while satisfying the target E-field magnitude constraint Eq. 3. We implemented the minimization of Eq. 10 in `pynibs.optimization.foc_opt()` with the use of SciPy (Virtanen et al., 2020) and the BFGS solver. Note that this formulation does not yet take into account the current rate constraints (see next section).

### Current-rate constraint

Besides ensuring the target E-field magnitude, real world scenarios also need to take into account the maximum achievable current rate, which is limited by the design of the stimulator electronics, the mechanical and thermal constraints of the coils, and parameters such as the pulse shape and stimulation frequency. Here, we utilize a maximal current rate in multicoil systems is *J*_*limit*_ = 100 A/µs, which is a typical value.

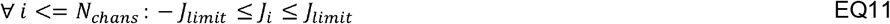

### Quality Metrics

We quantify focality by the ratio between squared E-field magnitudes at the target and at off-targets over the entire cortex:

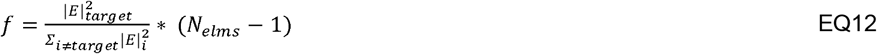

Besides this generic definition of focality we introduce two additional metrics to assess the realized cortical E-field patterns with respect to practically relevant parameters: *Target2Max* and *OverstimulatedArea*:

*Target2Max* (T2M, 0 < *T*2*M* ≤ 1) quantifies the E-field magnitude at the stimulation target in relation to the maximum across the whole cortex:

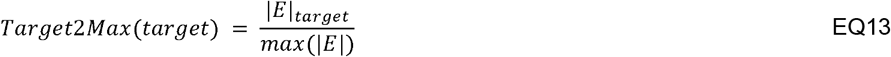

If the target receives the highest E-field magnitude, *Target2Max* equals one, while *Target2Max<1* indicates that the required E-field magnitude is higher elsewhere.

*OverstimulatedArea* quantifies the part of the cortex that is stimulated above a certain fraction *p* (0 < *p* < 1) of the target E-field magnitude (e.g., *p*=0.9 for more than 90% of the target magnitude).

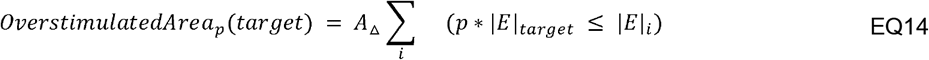

Here, *A*_Δ_ is the average element surface area and |*E*|_*i*_ is the E-field strength of element *i* in the rest of the cortex. Thus, *OA*_*p*_ quantifies the spread of the E-field w.r.t. a fraction *p* of with . equals the half-value area (Deng et al., 2013). *OverstimulatedArea* can also be expressed as a relative metric (), if is replaced by, with being the number of elements in the cortex.

It should be noted that *Target2Max* and *OverstimulatedArea* may have multiple, non-unique minima for different sets of input currents, as *Target2Max* saturates at 1 and *OverstimulatedArea* at 0. See SI-Quality metrics for E-field examples of cortical targets with different stimulation profiles for spherical (Figs. S5 and S6) and realistic head models (Figs. S7 and S8).

### Focality vs. target E-field magnitude

The optimal current rate solution as a function of the desired target E-field magnitude follows a scaled version of the focal solution, where the quality metrics (*f, Target2Max, OverstimulatedArea*) remain unchanged (see Fig. 3). Above a certain target strength, where the current rate constraint is violated (grey dotted line in Fig. 3), starts deviating from towards, and the quality metrics deteriorate. Above the maximal target E-field strength no solution exists.

**Figure 3.**
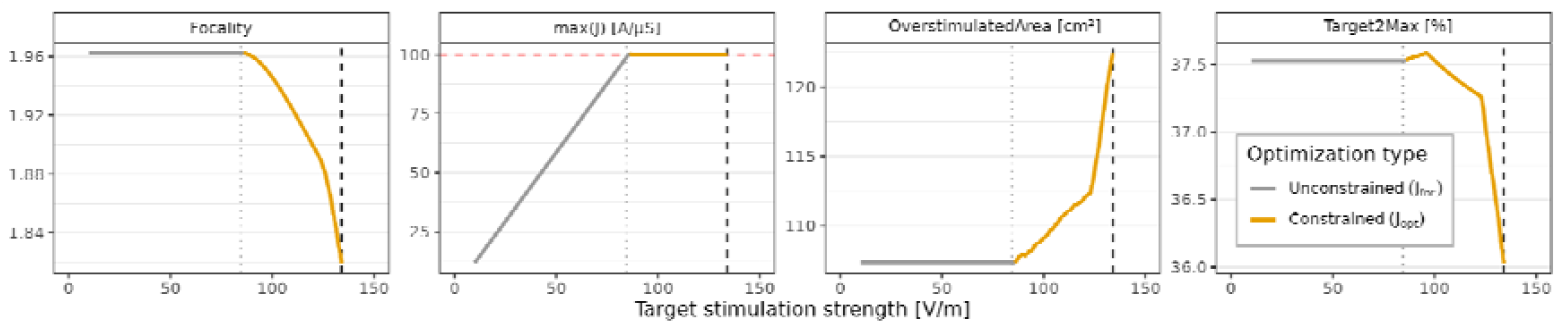
Stimulation strength effects on focality. Focality, maximum current rates, *OverstimulatedArea*, and *Target2Max* metrics, all plotted against the required E-field magnitude in one target. As long as the solution of Eq. 10 (*J*_*foc*_) does not violate the current rate constraints, *OverstimulatedArea*, and *Target2Max* remain constant, while the maximum current increases linearly with the required E-field magnitude (grey line), cf. Fig. 2a-b. For higher target E-field magnitudes (Fig. 2c), the constrained optimization (*J*_*opt*_) yields less focal stimulation (yellow line). Dashed red line: current-rate constraint (100 A/µs). Dashed grey line: Maximum target E-field magnitude feasible at this cortical target. Dotted line: maximum unconstrained stimulation.

## Results

### Effects of the region-of-interest definition on focality optimization

For any cortical target, the focality optimization identifies the channel-wise current rates that minimize off-target stimulation within a predefined ROI, while staying within the allowed current rate range. The focality optimization thus depends on the selection of the ROI. Its size, shape, and position with respect to the coil array placement affect the solution in terms of the achieved focality and of the resulting current rate vector. This is due to ROI-based leadfield matrices *L* that enter Eq. 1 and Eq. 10, forcing E-field suppression throughout the ROI.

In order to identify effects of the ROI definition on the optimization and to derive guidelines for ROI definition, we performed the focality optimization for all targets in three circular ROIs below the mTMS coil arrays in realistic head models and for a whole brain ROI in a spherical head model. See Fig. 4 for an exemplary subject with circular ROIs of different sizes contrasting the three different metrics *focality, OverstimulatedArea*, and *Target2Max* for the planar mTMS system, Figs. S1-S3 for details across the mTMS systems, and Fig S4 for the spherical head model results. Across all quality metrics, the optimization performs best for superficial (i.e., gyral) targets as compared to deeper targets. For targets very deep in the sulci, up to 120 cm^2^ receive more than 90% of the target strength (i.e., >90 V/m for 100 V/m target stimulation), and the largest off-target E-fields amount to twice the target E-field (e.g. 200 V/m). Moreover, the quality of the optimization depends on the size of the ROI, with medium-to-large ROIs yielding best suppression of off-target activity, with whole-brain ROIs not further improving focality (see Fig. S4).

**Figure 4.**
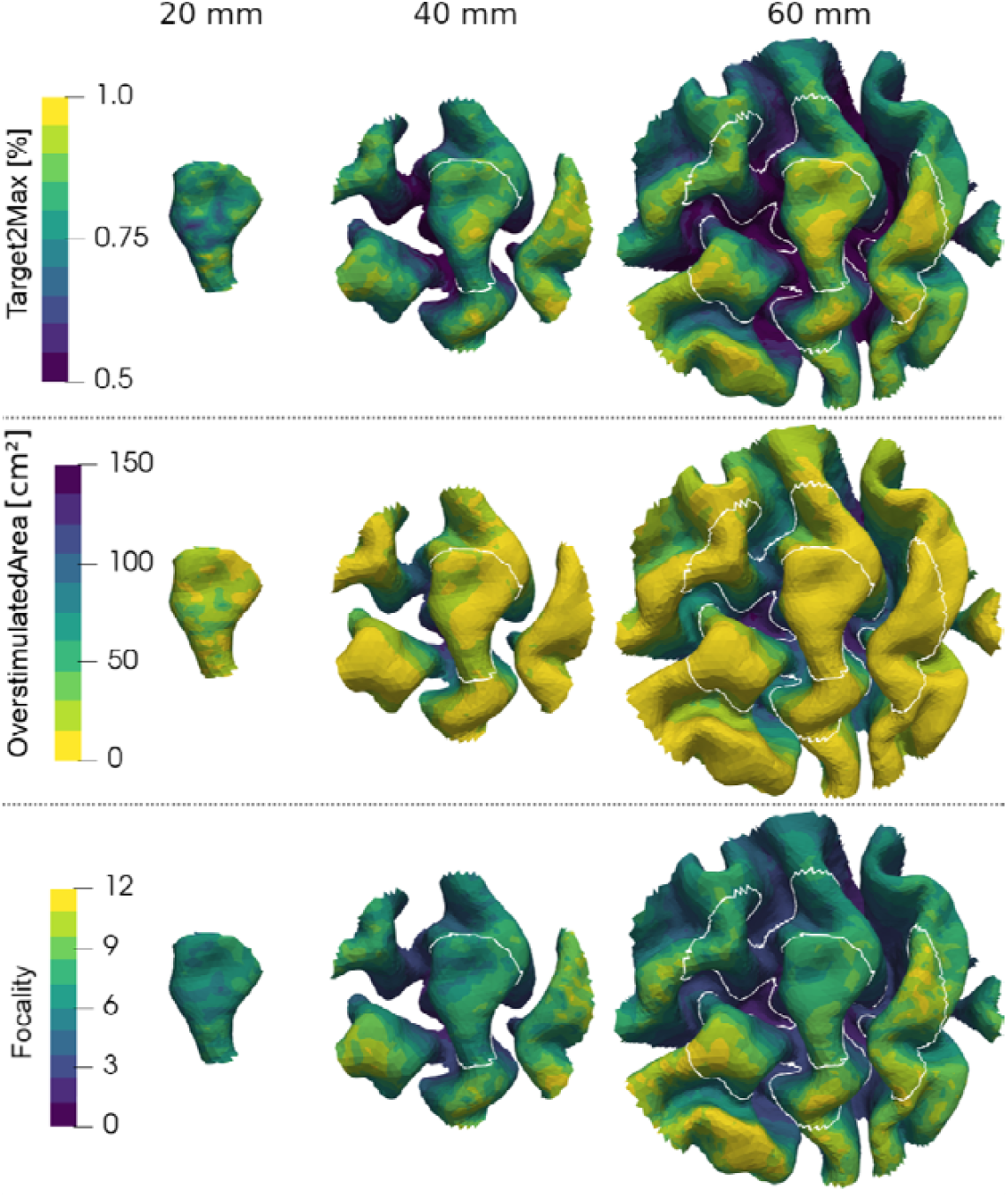
The ROI size affects the stimulation optimization. Shown for an exemplary subject with a 5-chan planar mTMS array and spherical grey matter ROI centered on the hand knob. For each element within each ROI, the optimal current rates were determined. Top row: *Target2Max* ratio, quantifying the degree of overstimulation. Middle row: *OverstimulatedArea*_*0*.*9*_, quantifying the area that receives higher stimulation than the target element. Bottom row: Stimulation focality quantified with (Eq. 10). Quality metrics were computed with respect to the entire cortex while current rate optimization is performed considering the ROI. Color scales: Yellow depicts better, blue depicts worse solutions. For small ROIs (left column), focality optimization performs poorly, yielding large overstimulated areas and low *Target2Max* values— indicating unfocal stimulation with stimulation hotspots outside of the ROI. Larger ROIs (middle and right columns) lead to better solutions, especially on the gyral crowns as more cortical elements are included in the optimization. White lines: small and medium ROI outlines. See Figs. S1-S3 for the other mTMS systems.

Together, the different mTMS systems with their specific coil geometries render a strategic placement of the coil arrays necessary, depending on the set of stimulation targets. The size of the ROI further affects the optimization, with a very small ROI yielding suboptimal stimulation focality. From here on, we utilize a left-hemispheric primary motor ROI (M1, S1, PMd; Numssen et al., 2021) as an example of a functionally defined and widely examined brain region for TMS studies (Jing et al., 2025).

### Effects of mTMS geometry on focality optimization

To identify effects of the mTMS coil geometry on the focality optimization we optimized the current rates for each element in a primary motor cortex ROI individually. This was performed for nine subjects, yielding about 12500 targets, and thus, optimizations, per subject for one placement per mTMS system (c.f. Fig. 1c). For this setup, all targets could be stimulated with at least 100 V/m at 100 A/µs, i.e. no targets needed to be excluded from the analyses (Table S1). The minimum achievable stimulation strength differs across mTMS designs: for the 12-chan spherical system, each location within the ROI could be stimulated with at least 223 V/m, in contrast to about 130 V/m for the 5 and 6 channel systems. In terms of the highest stimulation intensity that can be achieved anywhere, the planar design slightly outperformed the spherical ones with almost 600 V/m.

Accordingly, for each target, the focality optimization was either not affected by the current constraint, for example for superficial targets (*J*_*foc*_ = *J*_*opt*_, c.f. Fig. 2a, 2b), or it was affected (*J*_*foc*_ ≠*J*_*opt*_, Fig. 2c). For the 5-chan planar mTMS system, most targets in the ROI coil could be stimulated without taking the current constraint into account, while the 6-channel, and to an even stronger degree, the 12-channel systems required constraint-aware optimization (Fig. 5, left column).

**Figure 5.**
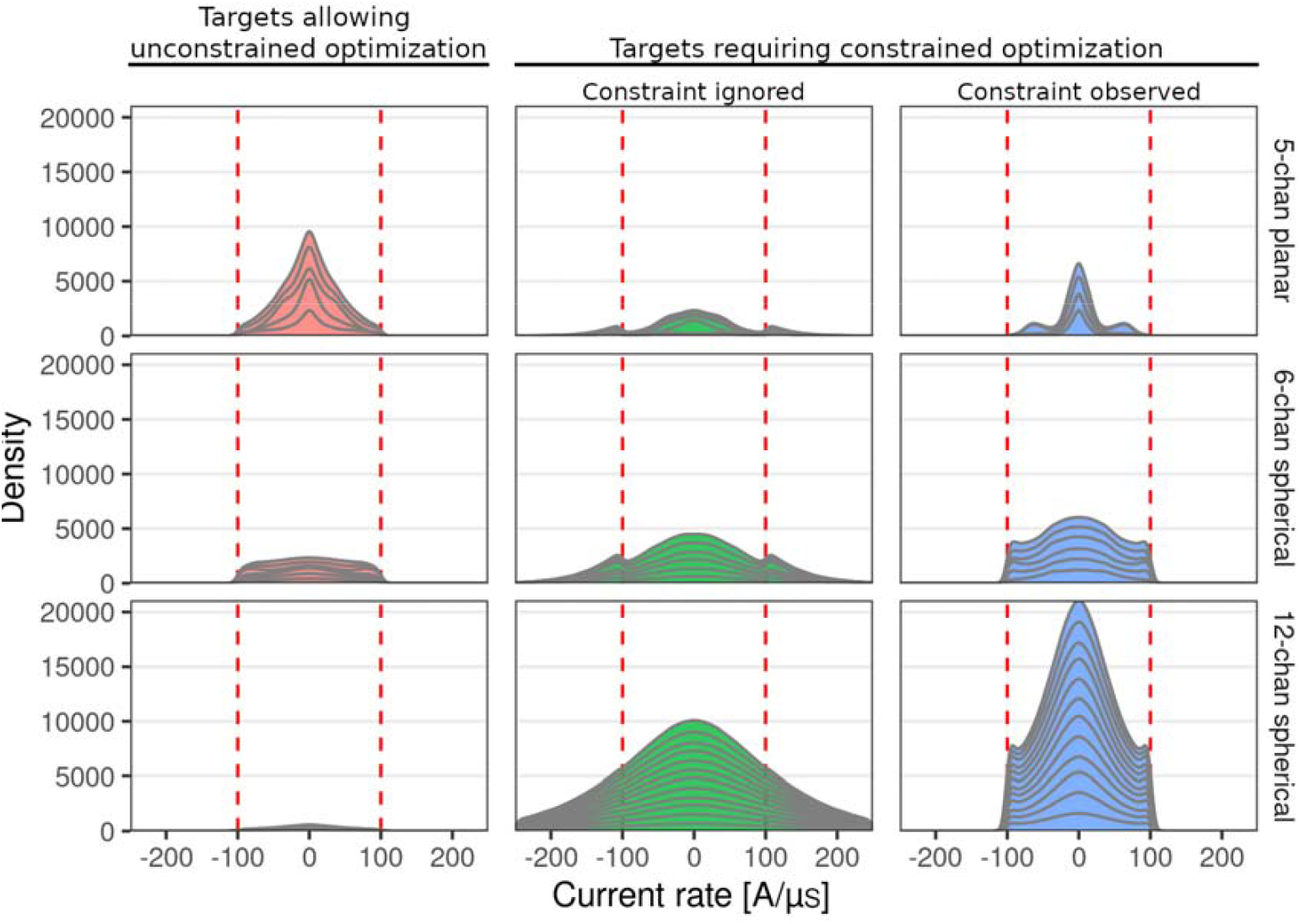
Coil array geometries affect selection of current rates. The optimized, single-channel current rates are shown for focality optimized stimulation of each target in the motor cortices of 9 subjects (∼12500 targets per subject; one coil array placement) for 100 V/m target strength and 100 A/µs current rate constraint (dashed vertical lines). Left: Subsample of cortical targets that did not require the current constraint to yield focal stimulation at 100 V/m. Center and right columns: Subsample of targets that did require the constraint optimization. For these, results are shown for the case that the current-constraints are ignored (green) or observed (blue). Top: planar 5 channel mTMS coil; Middle: 6-channel spherical TMS coil (middle); Bottom: 12 channel spherical mTMS coil (bottom). Grey lines: single-channels.

For targets affected by the current constraint, we performed an unconstrained optimization and a constrained one to provide insights into the optimization (Fig. 5, middle and right columns). Without constraint, the focality optimization selected maximum current rates of 326.91 A/µs, 306.67 A/µs, and 692.20 A/µs for the 5-chan planar, 6-chan spherical, and 12-chan spherical mTMS systems, respectively, identifying considerably stronger constraint-effects for the 12-channel spherical system compared to 5-channel and 6-channel mTMS arrays. Analyzing the channel-wise effects of the constrained optimization (Figs. S9-S11) identified strong differences between coil systems. For the 5-channel planar system, enforcing the constraint shifted current-rates from the cloverleaf and round coils towards both figure-of-eight coils (Fig. S9). For the 6-channel spherical design, the x- and y-channels receive high levels of current rates, most probably due to their increased distance to the cortical surface (Fig. S10). The 12-channel spherical design shows, in contrast, equal recruitment of all channels, irrespective of their spatial orientation (Fig. S11).

In summary, all three mTMS designs allow effective stimulation of all ROI targets, but a higher number of mTMS channels leads to more cases and stronger intervention of the constrained optimization. Furthermore, single-channel current-rate analysis identifies higher current-rates for the lower figure-of-eight channel of the 5-channel planar mTMS system and, more critical due to the higher number of occurrences, high demands on the x- and y-channels of the 6-channel spherical mTMS system.

### Multi-channel versus single-channel

To critically test if mTMS systems can effectively stimulate multiple cortical targets without repositioning of the coil array, we 1) optimized the mTMS placement for the center of the ROI, i.e. the handknob gyral crown, and 2) focality-optimized stimulation for each cortical target, i.e. each element in the ROI. For both sTMS coils, in contrast, we optimized placements for each cortical target. While this approach underestimates the maximum focality of the mTMS systems (which would require repositioning the coil array each time), it allows for a pragmatic assessment of how mTMS can be utilized in real life.

Figure 6 summarizes the quality measures for all target locations in the motor cortices of 9 subjects. When comparing sTMS and mTMS, we observe that a freely movable figure-of-eight coil often allows higher focality than a stationary mTMS array, especially for superficial targets (Fig. 6, top row). The focality metric (Eq. 3) quantifies the average stimulation at all off-target locations in relation to the field strength at the target. However, when considering metrics that selectively emphasize more substantial overstimulation in off-target areas, such as stimulation higher than in the target (*Target2Max*) or above a certain level (*OverstimulatedArea*), the mTMS arrays yield comparable results (Fig. 6, middle and bottom row).

**Figure 6.**
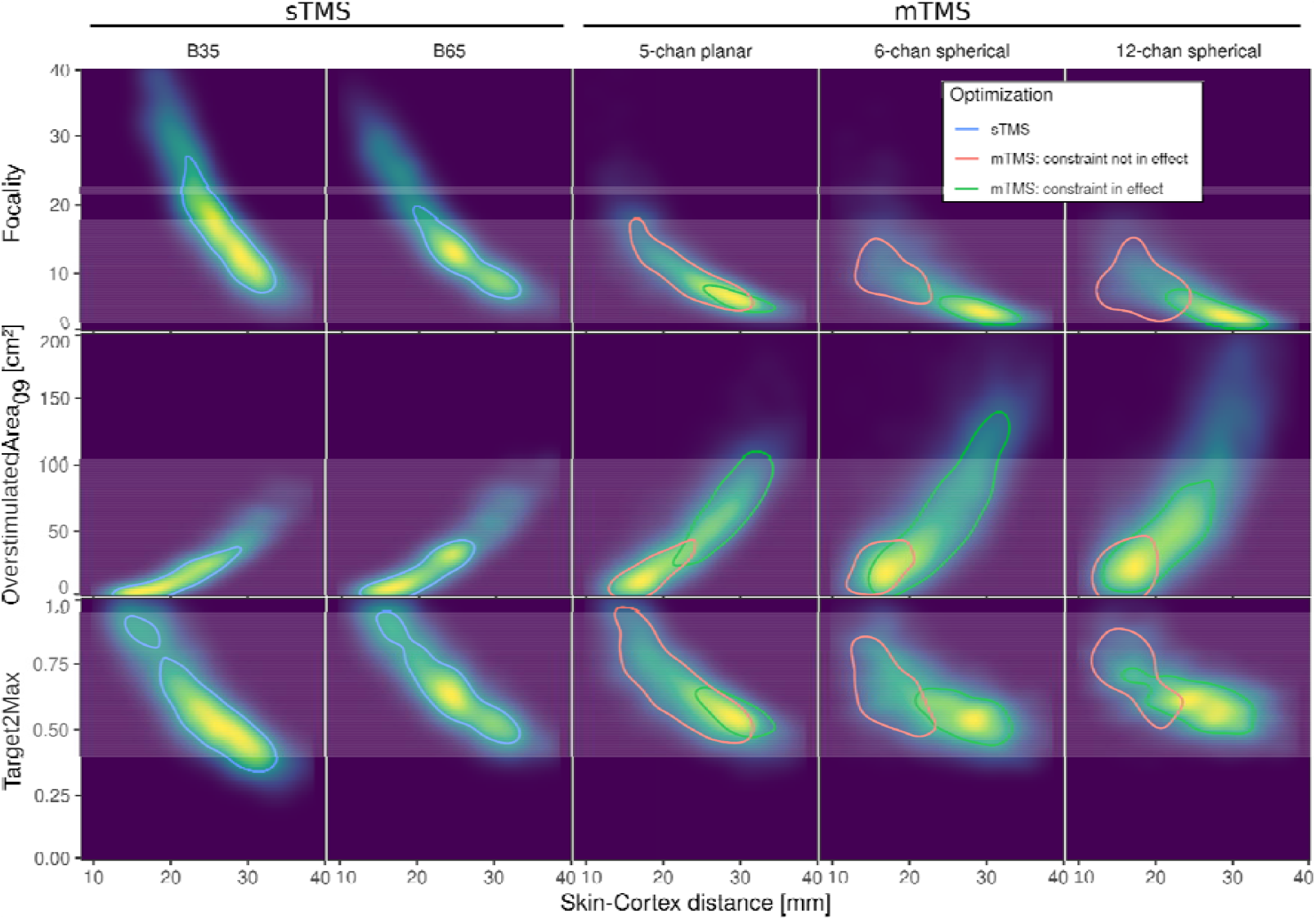
mTMS and sTMS systems yield comparable performances. Stimulation optimization results for 1000 targets in the motor cortex of nine subjects are shown. For sTMS, the coil placement was optimized for each target, for mTMS the coil array placement was optimized for the center of the ROI. Top: Focality versus target depth (skin-cortex distance). Deeper targets result in less focal stimulation, with sTMS yielding higher focality at superficial target locations. Middle: *OverstimulatedArea*_*09*_ versus target depth. sTMS and mTMS systems perform similarly, with the 6-channel and 12-channel spherical systems performing slightly worse for deeper targets. Note that lower values mean better stimulation. Bottom: The 5-channel planar system yields similar *Target2Max* performance as the sTMS coil across all target depths. The spherical systems perform slightly worse. Note that here higher values mean better performance. Overall, the mTMS systems (right) show slightly worse, but comparable performance as the sTMS coils (left). For the mTMS systems, more channels lead to more complex optimization, i.e. more cases of the constrained optimization (green vs. red contour). Contours delineate 50% of the data per condition; red for mTMS without current rate constraint becoming effective, green for mTMS with current rate constraint becoming effective, blue for |E|-optimal sTMS coil placement.

Among the mTMS systems, the planar design performs slightly better in terms of focality and OverstimulatedArea, especially for deeper targets. Within the spherical designs, more channels improve all three metrics.

In general, the mTMS systems perform surprisingly well compared to the sTMS coils, taking into account that the sTMS coils were repositioned for each target, while the mTMS coils were placed over the center of the ROI for all target stimulations.

## Discussion

Multi□channel TMS (mTMS) promises to overcome key limitations of traditional single□channel TMS (sTMS). By electronically shaping the induced E□field, mTMS enables stimulation of several cortical targets in rapid succession without physically repositioning the coil. This capability has the potential to revolutionize paradigms such as paired□pulse stimulation and to support automated, closed□loop localization of stimulation targets (Tervo et□al.,□2022; Matsuda et□al.,□2025; Granö et□al.,□2025a). Achieving this, however, critically depends on stimulation focality: the higher the focality, the finer the spatial resolution with which cortical targets can be addressed. In this work, we introduce a rigorous optimization framework that identifies the global minima of current sets to maximize focality under realistic hardware constraints in a very efficient manner. We further provide evaluation metrics that enable systematic comparison of mTMS and sTMS designs on individualized head models. Together, these contributions not only quantify how mTMS designs perform relative to established sTMS coils, but also provide a practical toolset for tailoring mTMS stimulation in future experimental and clinical applications and compare performance of different mTMS geometries.

### Quality assessment

To assess stimulation quality in a straightforward and practically applicable manner, we employed three complementary metrics.

The *focality* metric (Eq. 3) quantifies the ratio between the E□field strength at the target and the average E□field strength across off□target locations. While similar to the optimization goal function, this metric considers off□target locations throughout the entire cortex rather than restricting the evaluation to the ROI. By design, it is sensitive not only to strongly stimulated off□target areas but also to widespread regions of low□level stimulation.

The *OverstimulatedArea*_*X%*_ metric quantifies the proportion of cortical surface that is stimulated with an E□field magnitude equal to or exceeding *X*□% of the target’s stimulation strength. This metric specifically captures off-target regions where the induced field is likely to be physiologically effective (for example, ≥□90□% of the target E-field), while disregarding regions with stimulation levels too weak to trigger neuronal activation. Thus, *OverstimulatedArea*_*X*%_ does not penalize harmless low□level stimulation. Importantly, this metric requires a careful choice of the threshold *X*.

Finally, the *Target2Max* metric captures the maximum degree of overstimulation across all off□target regions. It indicates, for example, how much unintended stimulation is required in superficial areas to achieve the desired E□field strength at a deeper target. While these three metrics are interrelated, each emphasizes a different aspect of off□target stimulation. As the focality goal function minimizes the off-target stimulation strength while keeping the target stimulation constant, the presented results for *OverstimulatedArea* and *Target2Max* do not necessarily represent the minimal possible values for the investigated sTMS and mTMS coil arrays. Future research could implement limits of both quality metrics as additional constraints into the focality optimization framework to limit, e.g. the ratio of off-target stimulation to a specific *Target2Max* threshold. Taken together, they provide a comprehensive and meaningful characterization of the stimulation profile of a TMS system.

### mTMS geometries

We applied the focality framework to three mTMS systems with distinct geometries: a 5□channel planar mTMS transducer (Koponen et□al.,□2018) and two spherical transducers incorporating 2 and 4 three□channel, three□axis units, resulting in 6 and 12 channels, respectively (Navarro de Lara et al., 2021). Overall, these systems show comparable performance in terms of stimulation focality. The planar mTMS system replicated sTMS figure-of-eight focality, in line with its design objectives. For the spherical system, increasing the number of channels provides a slight improvement in focality. However, it also increases the complexity of the optimization process, as a larger proportion of cortical targets requires constrained optimization to keep current rates within safe operational limits.

Integrating a realistic current□rate constraint into the optimization is crucial - not only for safe mTMS operation but also for a fair assessment of system performance. Higher coil currents impose greater physical strain on the hardware and lead to increased heat dissipation. This becomes particularly critical when mTMS is extended from single□pulse to repetitive stimulation, where thermal management will be a central engineering challenge. Assessing this parameter early in the development process is therefore essential. Our focality optimization framework supports detailed single□channel analyses of required current rates, enabling the identification of potential bottlenecks already at the coil design stage.

### mTMS comparison with sTMS

We critically assessed the mTMS against two widely used, commercially available figure□of□eight sTMS coils. Importantly, the mTMS coils were positioned only once, above the center of the ROI, whereas the sTMS coils were repositioned for each cortical target to achieve optimal stimulation. This design directly tested whether mTMS systems can deliver on their promise of stimulating multiple targets in succession without physical repositioning, while achieving focality comparable to standard sTMS approaches.

For superficial targets, the ability to reposition the sTMS coils resulted in slightly better stimulation focality. However, for medium and deeper cortical targets (>25 mm SCD) mTMS and sTMS systems perform very similarly. This strong performance demonstrates that mTMS systems can provide equal stimulation quality, while offering the practical advantage of electronic targeting. In other words, mTMS not only matches sTMS in many focality benchmarks but also enables efficient multi□target stimulation without repeated coil adjustments, paving the way for more flexible experimental designs and novel therapeutic protocols.

### Outlook

These results highlight that assessments of mTMS coil array performance need to be performed for specific use-cases, supported by suitable metrics. This includes not only the inclusion of further constraints or penalties into the goal function (e.g., versions of the quality metrics introduced in this manuscript), but also the specific choice of stimulation parameters, such as the cortical target region, target stimulation strength, and stimulator limits, into the assessment. Importantly, our results show significant impact not only of the coil geometries, but also of the head and brain morphologies. While this study utilized a realistic sample of healthy individuals across the lifespan (aged from 18 to 87 years) to provide a valid basis for analyses, future research should address the impact of subject factors such as age, sex (Hoornweder et al., 2024) or ethnicity (Zhang et al., 2025) on stimulation focality more thoroughly by using larger samples. Because focality optimization and mTMS field steering rely on accurate, subject-specific E⍰field estimation, patients with altered brain structure—such as those with stroke lesions, cortical malformations, or prior resections (Minjoli et al., 2017; Mantell et al. 2021)—stand to benefit most from further research on lesion⍰ and malformation⍰informed E⍰field modeling and tissue-condictivity individualization.

While mTMS is in an early stage, its implications for neuromodulation applications are significant as precision neurostimulation is key to improve NIBS across research, diagnostic and therapeutic settings (Sinisalo et al., 2025a; Sinisalo et al., 2024b; Souza et al., 2025b). The ability to electronically select the stimulation targets without physical coil repositioning allows, in principle, for online optimization of target engagement (Lioumis et al., 2025; Tik et al., 2017), for example when combined with concurrent fMRI (Navarro de Lara et al., 2023; Souza et al., 2025b), EEG (Sinisalo et al., 2025b), or downstream outcome metrics, such as ECG (Feng, Martin et al., 2026). Alongside their application in motor cortex somatotopy mapping, focality-optimized mTMS pulses may also provide a means to investigate a newly proposed framework describing the complexity of motor cortex organization (Morishita et al., 2025). As a crucial next step, mTMS will need to support repetitive stimulation protocols to fully exploit its potential for therapeutic applications, for example intermixing facilitatory and inhibitory protocols targeting different cortical targets.

### Limitations

As the focality goal function minimizes *f*, i.e. the off-target stimulation strength while keeping the target stimulation constant, the presented results for *OverstimulatedArea* and *Target2Max* do not necessarily represent the global optima of these metrics for the respective coil arrays. While utilizing individual head and brain meshes to compute the mTMS lead fields takes the individual morphology into account during the focality optimization, current E-field simulation approaches ignore subject- or population-wise variability of tissue parameters, such as conductivity (Weise et al., 2025b) or grey-matter-density (Qi et al., 2025), both potentially affecting the TMS-induced E-field. Additionally, we specifically focus on the magnitude of the TMS-induced E-field, as currently no consensus exists regarding the directional sensitivity of the neuronal activation curve (e.g. Granö et al., 2025b; Zhao et al., 2026; Jing et al., 2025). Finally, the coil current limits and the specific coil models utilized in this study raise no claims of accurately representing reality but. Instead, they were chosen as a reasonable guess to present this focality framework. Especially for repetitive TMS, these will need to be revisited for future mTMS hardware generations (Sinisalo et al., 2025b).

## Conclusion

While mTMS aims to overcome fundamental limitations of sTMS - most notably the need for repeated physical repositioning – the increased flexibility of mTMS systems also brings increased complexity in their characterization and evaluation. Unlike sTMS coils, which have a fixed spatial profile, the focality of mTMS depends on many interacting factors, including the number and geometry of channels, their placement, and the current□rate configuration. Our analyses show that increasing the number of channels can modestly improve focality but also increases the need for constrained optimization to avoid exceeding safe current rates and to account for hardware limitations such as heat dissipation, especially in repetitive stimulation scenarios. Importantly, we demonstrated that even without repositioning, mTMS systems can achieve focality comparable to standard sTMS. Overall, mTMS performs remarkably well, highlighting its potential for efficient, multi□target stimulation.

With the focality framework and metrics presented here, we provide a practical and transparent approach for assessing mTMS performance under realistic constraints. We expect that these tools will support both developers and users of mTMS technology in systematically evaluating designs, identifying bottlenecks early in coil development, and enabling safer, more effective applications of mTMS in research and clinical settings.

## Data and Software Availability

The algorithms for the focality optimization and the focality-maximality balancing are made available through our Python package pyNIBS (v0.2026.1; Numssen et al., 2021). Scripts for meshing (using SimNIBS 4.5), ROI construction, lead field computation and optimization are accessible at the project’s GitLab repository https://gitlab.gwdg.de/tms-localization/papers/mtms_focality. The headmeshes constructed for nine individuals from the CamCAN dataset (Shafto et al., 2014) and the precomputed lead fields are available here: https://osf.io/7pumv/.

## Acknowledgement

We thank Victor Hugo Souza and Risto Ilmoniemi for providing information to model the 5-channel planar mTMS coil array and Mohammad Daneshzand, Evgenii Kim, and Aapo Nummenmaa for information on the 6-channel and 12-channel spherical mTMS models.

---

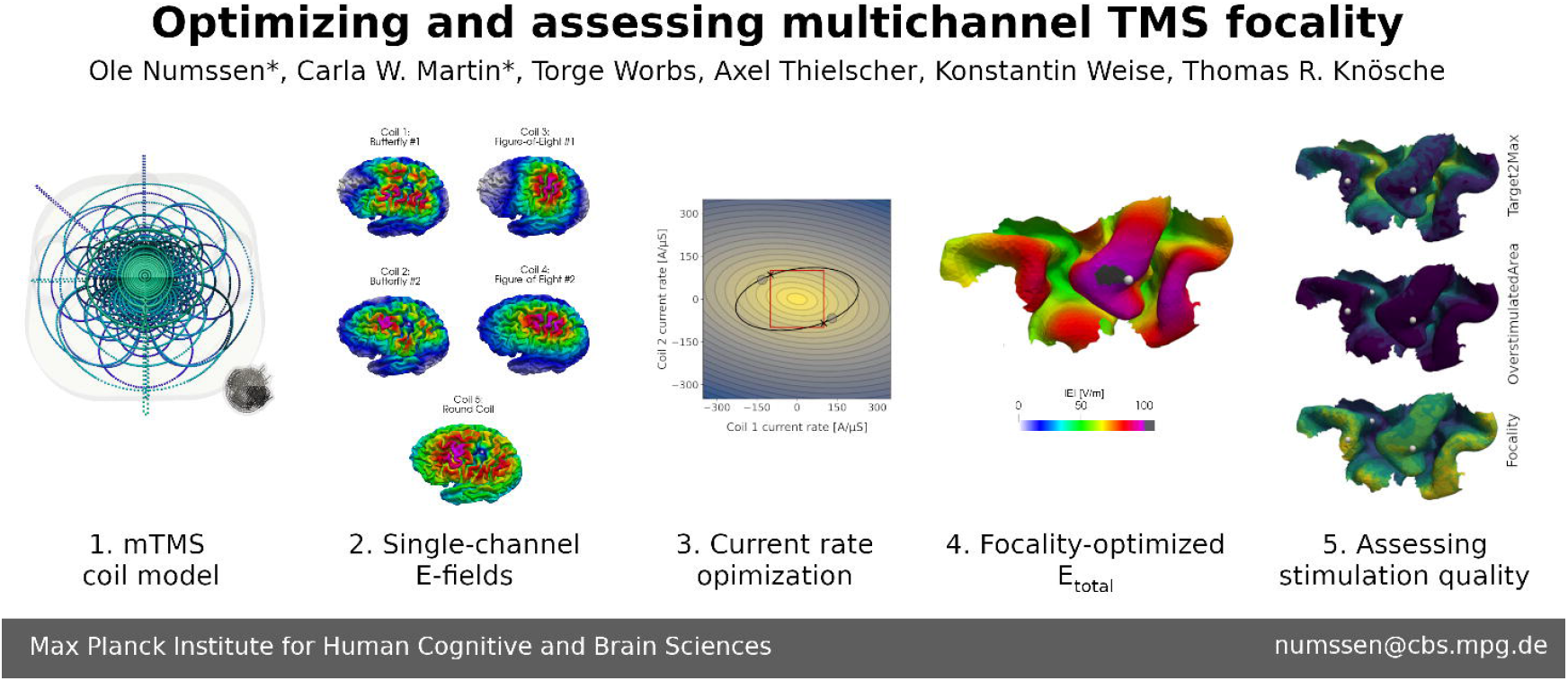

## Footnotes

1 There are always two solutions, because reversing all currents leads to the same values for Eq. 2 and Eq. 3.

