## Supplementary material for "Optimizing and assessing multichannel TMS focality": SI

Optimizing and assessing multichannel TMS focality - Supplemental Information

### Methods

#### Study Design

This study employed *in silico* experiments to optimize and evaluate the spatial focality of two major multi-channel TMS (mTMS) designs: a planar multi-coil design (Koponen et al., 2018) and a modular 3-axis design (Navarro de Lara et al, 2021). E-field simulations were performed using high-resolution individual head models derived from structural MRI data of ten healthy individuals across the lifespan. Spatial stimulation focality was optimized with a novel optimization algorithm that tunes coil currents to deliver a predefined E-field magnitude (stimulation strength) to a particular cortical target, while minimizing field spread at off-target locations and keeping the current rates in each coil within predefined limits. Two additional metrics were used to assess different aspects of stimulation focality: the *Target2Max* ratio, which measures the relative strength of stimulation at the target compared to the global peak, and the *Overstimulated Area*, which quantifies the cortical surface receiving stimulation above a specified relative threshold. The performance of the mTMS systems was compared against two different conventional single-channel figure-of-eight coils (sTMS). All scripts and coil models are made publicly available in a repository ([gitlab.gwdg.de/tms-localization/papers/mtms_focopt](https://gitlab.gwdg.de/tms-localization/papers/mtms_focopt)), all optimization algorithms and focality metrics in our Python package pyNIBS (v0.2025.8; [pynibs.readthedocs.io](https://pynibs.readthedocs.io/)), and the precomputed lead fields in an osf.io data repository (<https://osf.io/7pumv/>).

##### Head and brain meshes

Head and brain meshes were constructed for 9 individuals (5 female) from the CamCAN dataset (Shafto et al., 2014) spanning the adult lifespan (mean±SD age: 53.1±26.80) using SimNIBS 4.5 (Thielscher et al., 2015) from high-resolution T1- and T2-weighted MRI scans (1 mm³ voxel size). Tissue segmentation was performed using CHARM (Puonti et al., 2020) and FreeSurfer (Fischl, 2012), which identifies 15 distinct tissue types, including gray matter, white matter, cerebrospinal fluid (CSF), skull, and scalp.
For each individual, the whole brain grey matter mid-layer was extracted and used for all subsequent calculations, minimizing boundary artifacts and providing a consistent reference for evaluating focality metrics (pyNIBS; Numssen et al., 2021). Segmentation quality and cortical surface alignment were visually inspected to confirm anatomical accuracy.

##### TMS coil models

We built mTMS coil array models via line current elements with SimNIBS 4.5 (Worbs et al., 2025) based on publicly available information, to apply our focality framework to mTMS systems that are commercially available or currently under development. Three distinct mTMS coil array models were implemented to explore spatial focality and maximal stimulation strength: 1) The *5-chan planar* design consists of two four-leaf clover coils, two figure-of-eight coils, and one round coil, arranged in a stacked, layered configuration (Fig. 1a, left). 2) The *6-chan spherical* design features two 3-axis units, arranged to align with the scalp curvature with a 50 mm separation (Fig. 1a, center). Each unit consists of 3 orthogonally oriented round coils, allowing precise control over E-field direction and orientation. The modular nature of this design allows for the expansion to multiple units for broader cortical coverage. 3) The *12-chan spherical* design expands on the *6-chan spherical* configuration by incorporating two additional 3-axis coil units (Fig. 1a, right). Note that the coil array models used here are not intended to exactly represent any particular coil arrays used in past or future studies. Instead, they represent typical arrangements used so far to provide a testing ground for evaluating principal tradeoffs and limitations in mTMS coil arrays. The methodology provided in this paper can be easily applied to any other coil array by means of the provided scripts and example data sets. 4) For the purpose of comparison to sTMS, the MagVenture Cool-B35 and MCF-B65 coils (Drakaki et al., 2022) were selected as representative figure-of-eight coils, commonly used in TMS research and clinical applications due to their well-characterized stimulation profiles.

#### Coil Placement Optimization

To ensure optimal placement of the multi-channel (mTMS) and single-channel (sTMS) coils—both position and rotation on the skin surface—we performed a dedicated optimization before the focality analysis. For the sTMS coils, placement optimization is usually done by identifying the placement that maximizes the stimulation strength (∣E∣) at the cortical target. In contrast, mTMS coil placement optimization is less straightforward because the total induced field (E_total_) depends not only on coil placement but also on the current distribution across individual channels. To address this, we based the mTMS placement strategy on maximizing stimulation focality at the center of the region of interest (ROI). This approach provides system-specific placements and extrapolates the logic of the sTMS strategy while accounting for the unique designs of the different mTMS systems.
We used an adapted version of simnibs.tms_many_simulations() to compute single-channel E-fields for a number of candidate placements for each of the three mTMS systems. These candidate placements were centered above the cortical target on the skin surface (skin-coil distance set to 1 mm), with coil orientations varying from -90° to +90° in 5° steps, resulting in a total of 36 candidate placements. Each mTMS coil placement was adjusted using pynibs.shift_coil_to_cortex() to ensure that the minimum distance between any point on the coil housing and the skin surface was 1 mm for planar and bent coil array housings alike.
For sTMS coil placement optimization, we performed a standard approach, which maximizes |E| at the cortical target. Here, we chose the same optimization parameters (30 mm radius, 2 mm step size, 180° coil angle range, 7.5° step size) as for the mTMS placement optimizations. Importantly, for sTMS coils we optimized the placement for each cortical target in the ROI, whereas for mTMS we only optimized the placement for the center target.
For both approaches, we utilized the brute-force optimization implemented in SimNIBS (Weise, Numssen, et al., 2021) to compute lead fields for a large number of candidate coil positions and orientations (Nc = 3100) distributed systematically above the motor cortex. We decided to optimize the mTMS placements only for the center of the ROI while optimizing sTMS placements for each target location to critically test if mTMS does, indeed, allow to stimulate different cortical targets without coil replacements.

#### Spherical head model

We utilized a spherical head model from SimNIBS to perform the focality optimization without effects of individual morphology (Fig. S5). This head model consists of the same five major tissue types as realistic SimNIBS head models, with a radius of 95 mm and about 15 mm skin-cortex distance. We positioned the three mTMS coil arrays with about 2 mm distance between coil housing and skin surface. To mirror processing of realistic head models, we constructed a circular grey matter ROI at the grey-matter-CSF boundary, centered below the coil centers with a radius of 35 mm. For each target in the spherical grey matter ROI, we performed the focality optimization to maximize focality *f*; see Fig. S6 for element-wise focality results of all mTMS coil arrays*.* For the spherical ROI, the 5-chan planar and 12-chan spherical systems perform slightly better outside the ROI center, whereas the 6-chan spherical system yields best performances in the center of the ROI. We selected three targets (black triangles in Fig. S5) that show different performance to one another and across the mTMS systems to showcase their focality-optimized E-fields (Fig. S6). Target 1 is positioned at the center of the ROI, i.e. centered below the coil arrays. Target 2 is positioned outside of the ROI center, with good focality results for the 5-chan spherical array. Target 3 is positioned even further outside of the ROI center with good focality results for the 12-chan spherical system. Each target could be successfully stimulated with 100 V/m (pink color in Fig. S6) and the E-field hotspot is shifted towards each target. As expected, each individual E-field correlates with the quality metrics, such as larger areas of overstimulation (|E| > 100 V/m) for lower focality and vice versa.

#### Focality vs. target E-field magnitude

As detailed above, the optimal focality to stimulate a target with a particular E-field magnitude is determined by SI EQ10 when disregarding any current-rate constraints. For a target E-field threshold of $e=100 V/m$, we call the associated current-rate solution $J_{foc}$ (note that also ${-J}_{foc}$ is a solution). For different target E-field magnitudes, scaled versions of $J_{foc}$ apply. However, current-rate constraints may compromise, both, focality and E-field magnitude. As long as the required E-field in the target is sufficiently low, the focality optimization is not hampered by the current-rate constraint (Fig. 3). Hence, we obtain the maximum focality according to SI EQ2 and SI EQ3 (resp. SI EQ10) for this location through scaled versions of $J_{foc}$ (Fig. 2a and 2b). When increasing *e*, at some point, the optimal solutions start to violate the current-rate constraints (i.e., they are outside the hypercube) and the maximally achievable focality reduces. We call the current-rate solution that identifies the best focality while respecting the current-rate constraints $J_{opt}$ (Fig. 2c). If *e* is increased even further, no focality optimization is possible (Fig. 2d) anymore. Geometrically, this is equivalent to some of the corners of the hypercube being located on the ellipsoid-hypercylinder surface. This corresponds to the maximum E-field magnitude $e_{max}$ achievable for this target through the current rate vector $J_{opt} = J_{max}$, where each channel hits the maximum current rate $J_{limit}$ with different signs (i.e.,a corner of the hypercube). The achieved focality $f$ still varies between target locations and coil array placements, but is generally much lower than for $J_{foc}$. Beyond that point, the required target E-field magnitudes (see Fig. 2d) cannot be achieved.

### Results

#### Effects of the region-of-interest definition on focality optimization

Target2Max and OverstimulatedArea, which selectively emphasize relatively strong off-target E-fields, are affected by different ROI sizes while the focality metric, which shows the average suppression throughout the cortex, is barely affected. In contrast, a very small ROI size (20 mm diameter), potentially intriguing to use for computational efficiency, yields suboptimal results, as a large part of the stimulated cortex are not taken into account as off-targets, and therefore not actively suppressed.
For the 5-chan planar mTMS systems, 40 mm and 60 mm ROIs yield very similar focality results, whereas both spherical designs yield better performance for the 40 mm ROI (Figs. S1-S3). For the focality metric, the 6-chan spherical mTMS system yields best results at the center below the mTMS coil array, while the 5-chan planar and 12-chan spherical system yield better focality results outside of the center of the ROI (Fig. S2). For the OverstimulatedArea and Target2Max metrics, only the 12-channel spherical system yields better results outside of the center of the ROI, whereas the other two systems yield better performance at the ROI center, i.e., directly under the mTMS array (Fig. S1 and Fig. S3).

##### Target2Max

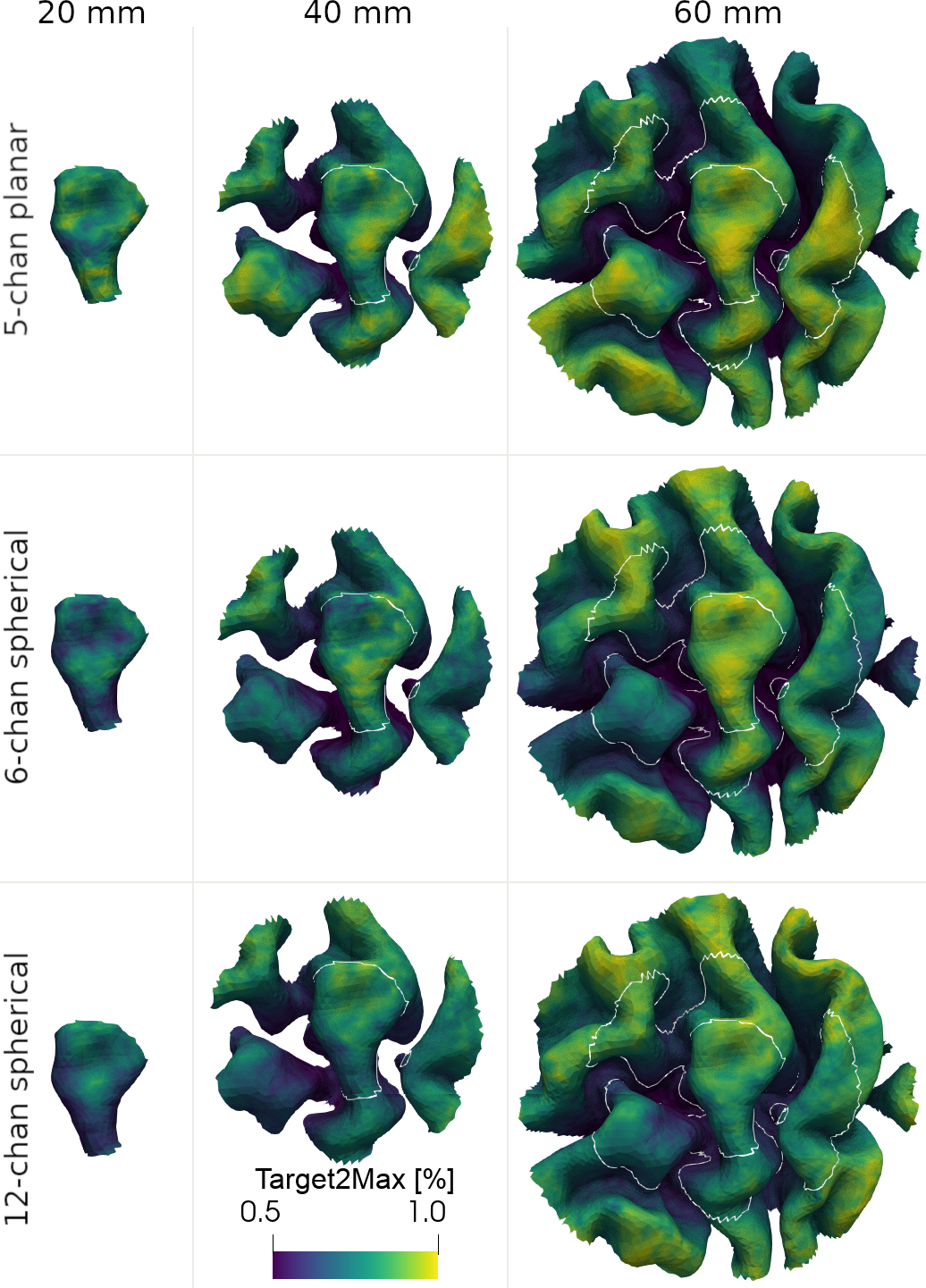

**Figure S1: Effect of ROI size on Target2Max.** Shown for an exemplary subject for a spherical grey matter ROI centered on the hand knob for three mTMS designs. For each element within the respective ROI, the optimal current rates were determined. Target2Max was computed with respect to the entire cortex. The yellow ends of the color scales mean better, the blue ones worse solutions. White lines: small and medium ROI outlines.

##### Focality

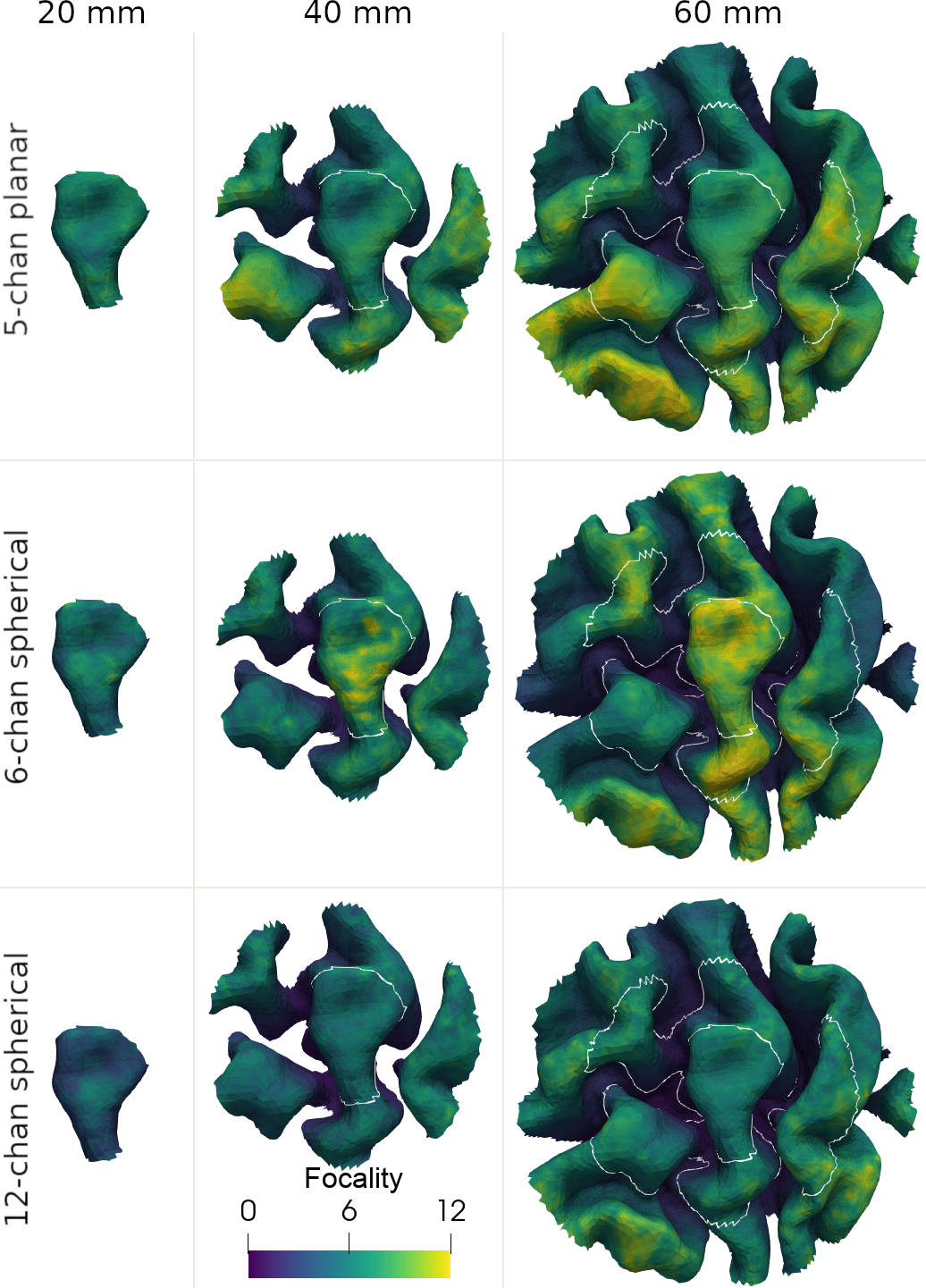

**Figure S2: Effect of ROI size on Focality.** Shown for an exemplary subject for a spherical grey matter ROI centered on the hand knob for three mTMS designs. For each element within the respective ROI, the optimal current rates were determined. Focaliy was computed with respect to the entire cortex. The yellow ends of the color scales mean better, the blue ones worse solutions. White lines: small and medium ROI outlines.

##### OverstimulatedArea

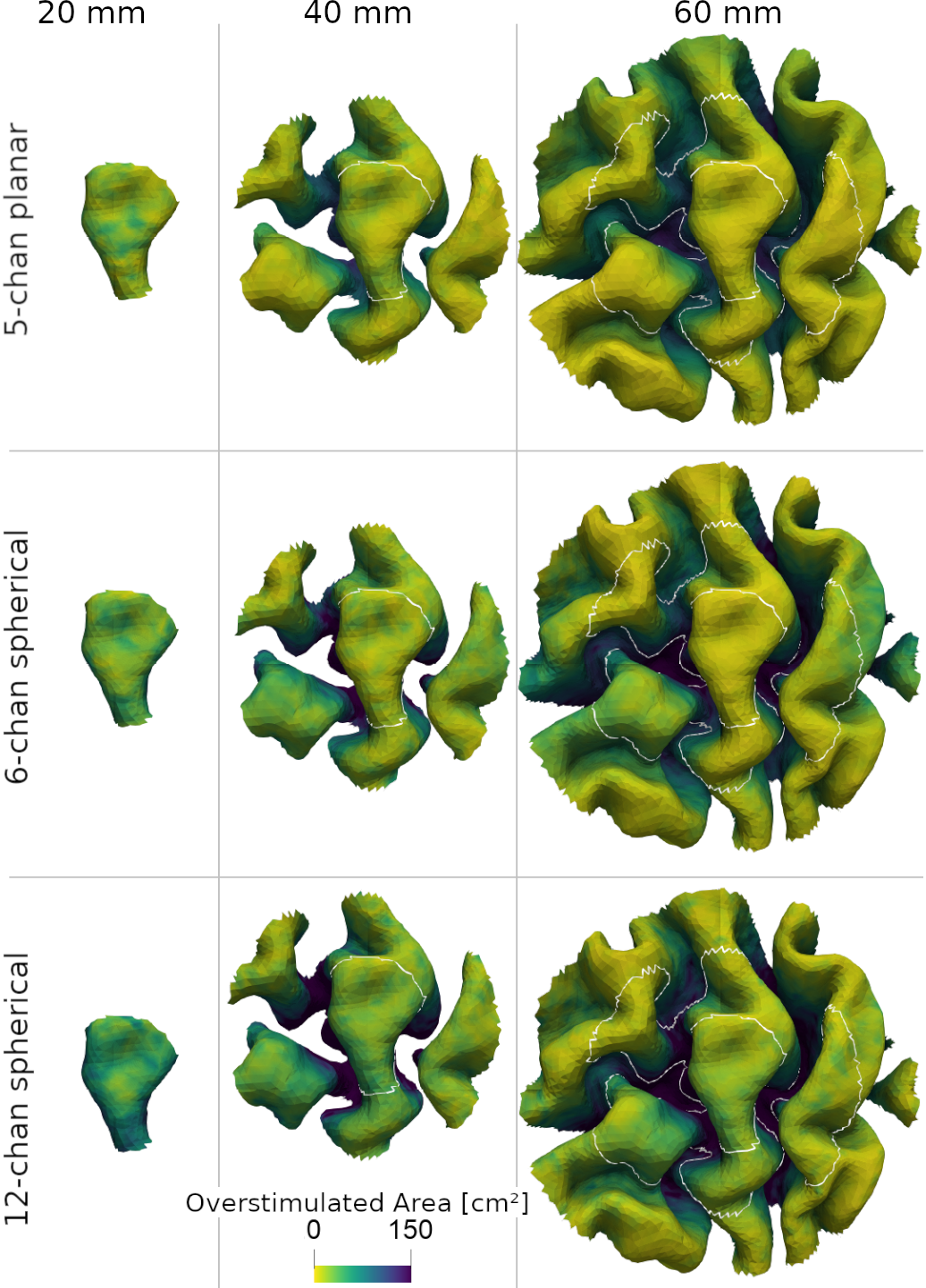

**Figure S3: Effect of ROI size on OverstimulatedArea.** Shown for an exemplary subject for a spherical grey matter ROI centered on the hand knob for three mTMS designs. For each element within the respective ROI, the optimal current rates were determined. OverstimulaedArea_0.9_ was computed with respect to the entire cortex. The yellow ends of the color scales mean better, the blue ones worse solutions. White lines: small and medium ROI outlines.

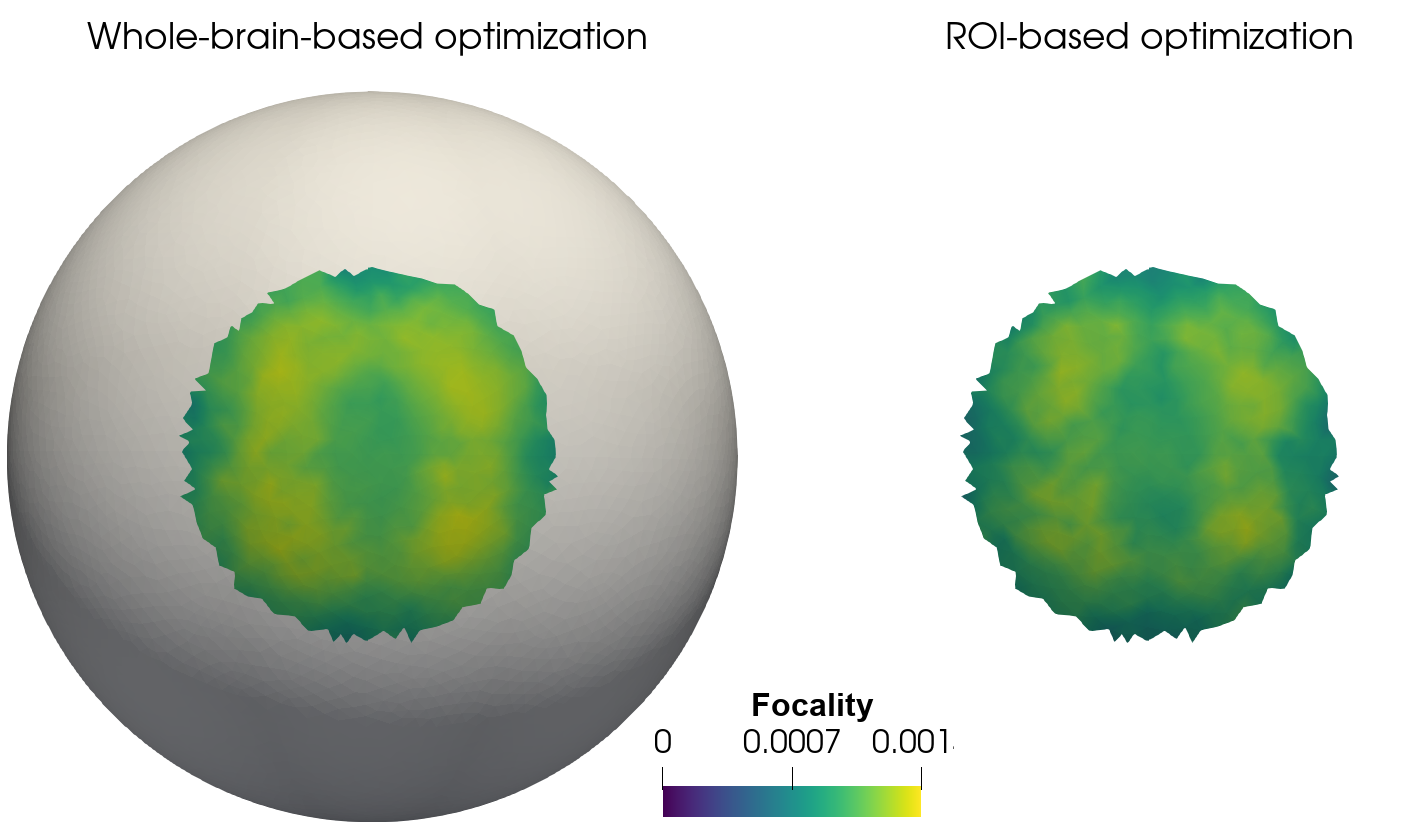

**Figure S4: Whole brain vs ROI focality optimization on spherical head model.** Increasing the ROI size to whole-brain grey matter surface does not further improve the focality optimization.

#### Stimulation focality

##### Spherical head model

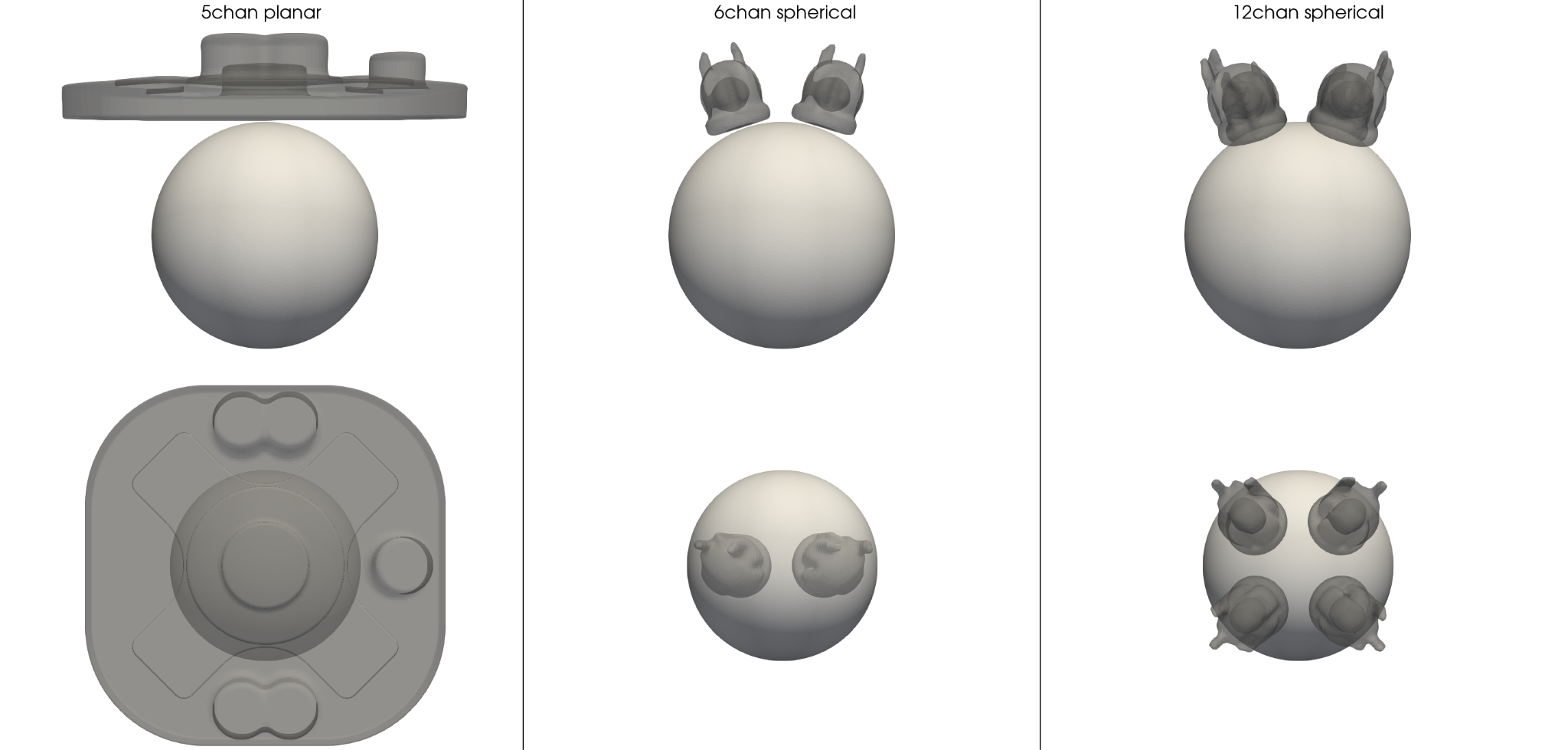

**Figure S5: mTMS coil array placements for the spherical head model.**

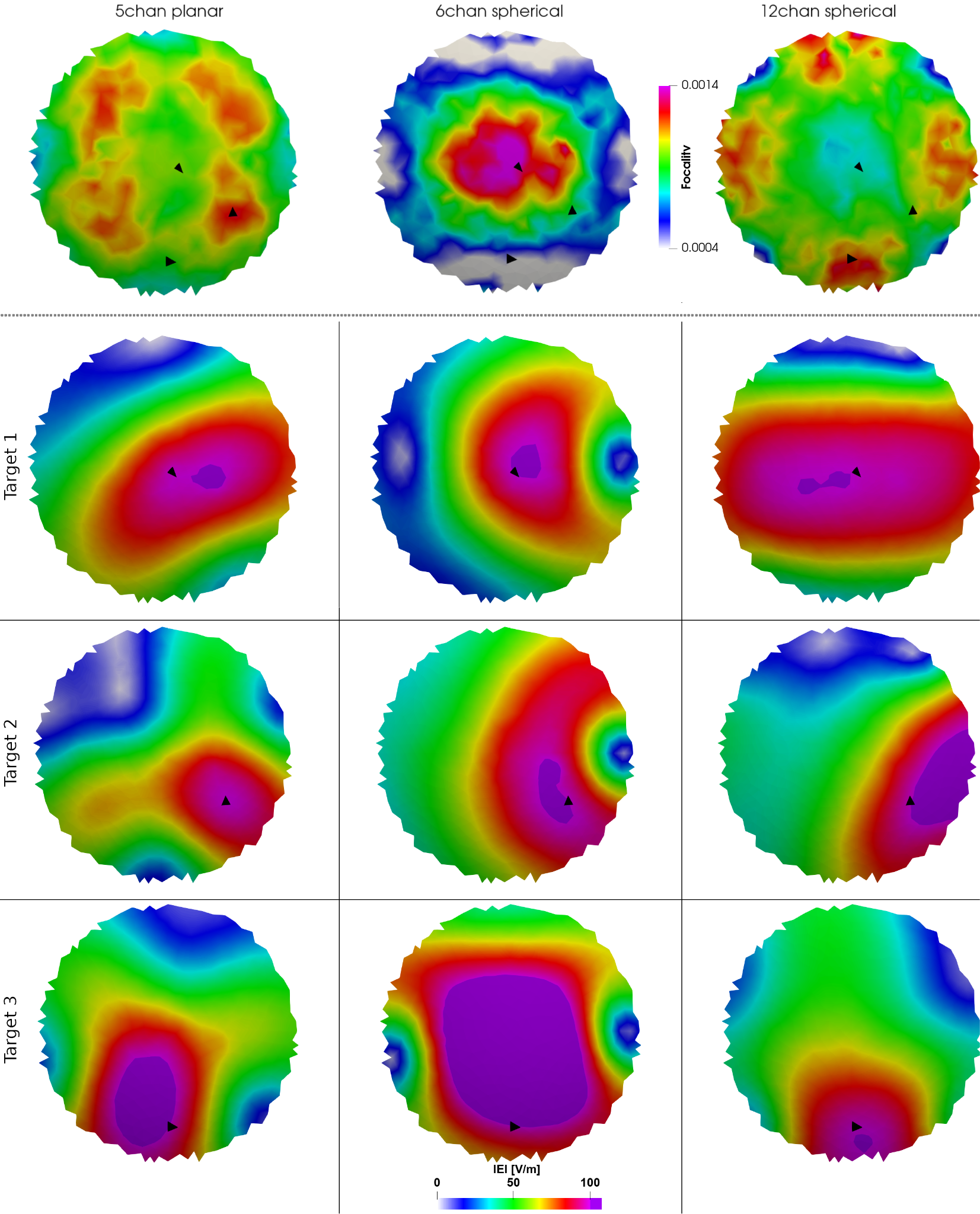

**Figure S6: Focality results in the spherical head model ROI.** For each target in the spherical ROI we performed the focality optimization for the 5-chan planar (left), 6-chan spherical (center) and 12-chan spherical mTMS coil array (right). Bottom: Focality-optimized E-fields for three exemplary targets (black triangles). Target 1: Centered below the mTMS coil array. Target 2 and Target 3: Targets outside of the ROI center. Color: E-field magnitude, cropped at 100 V/m.

##### Realistic head model

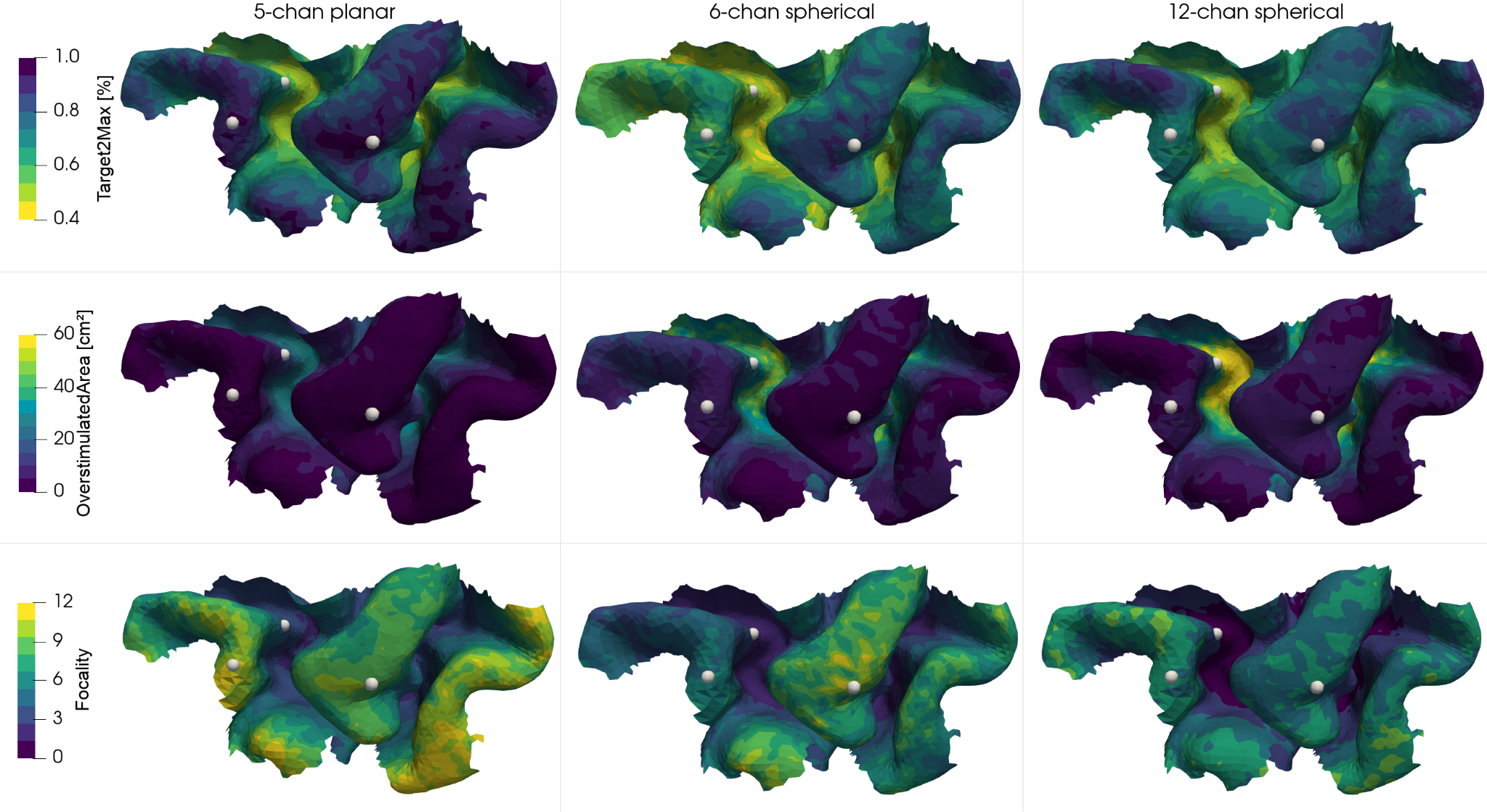

**Figure S7: Focality results in a realistic head model ROI.** Across mTMS coil arrays (columns) targets with low skin-cortex distance can be stimulated more focally than deeper targets. White spheres: Exemplary targets detailed in Fig. S7.

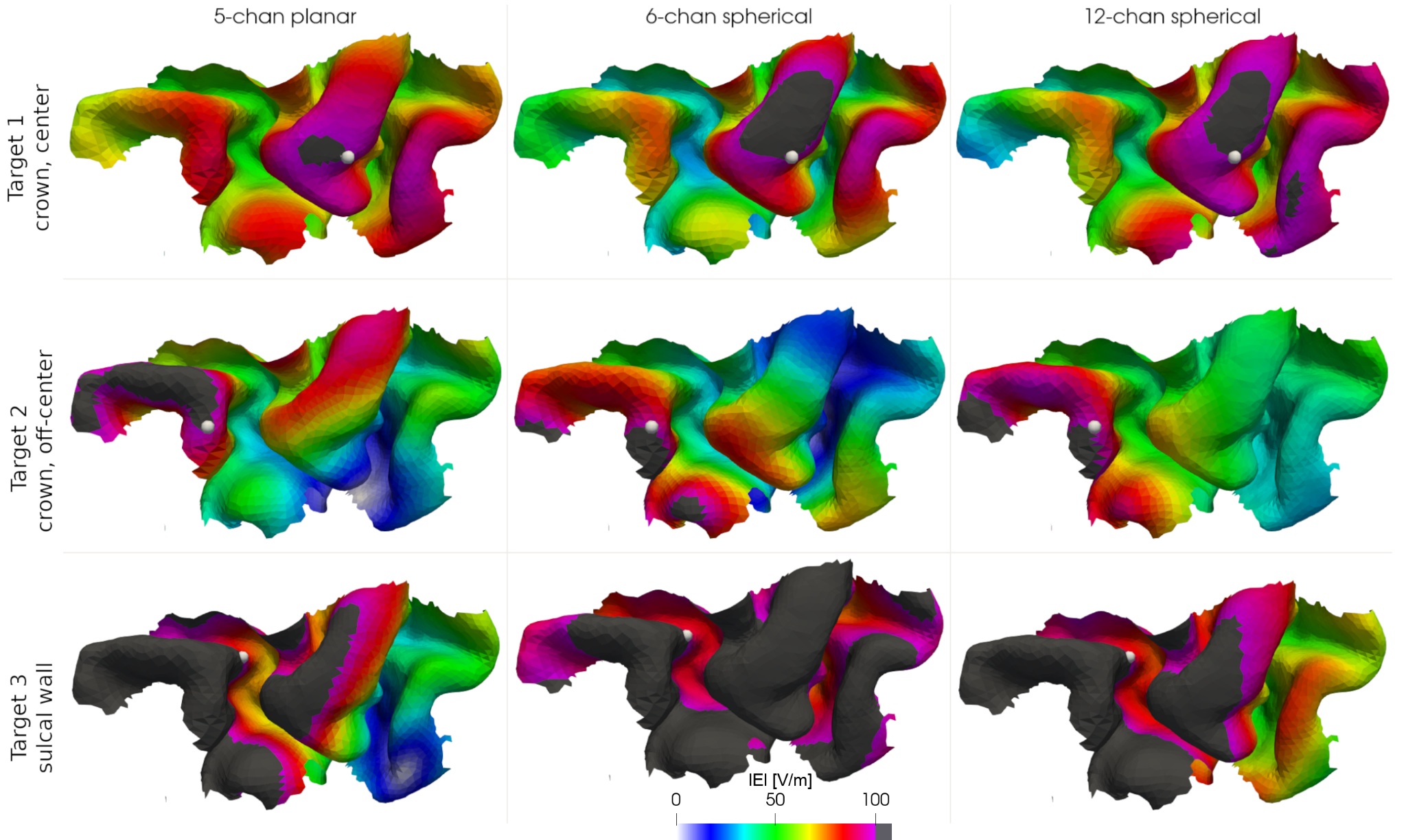

**Figure S8: Focality-optimized E-fields for three targets.** Across mTMS coil arrays (columns) gyral crown targets in the center of the ROI (Target 1) can be stimulated very focally. Targets outside of the ROI center show reduced focality (Target 2), while deeper targets (Target 3) yield significantly worse focality. Focality optimization was performed with 100 V/m at the target. White spheres: targets.

#### Effects of mTMS geometry on focality optimization

**Table S1**

*Maximum stimulation strength achievable across mTMS systems.*

| **mTMS system** | **min(\|E_max_\|) … max(\|E_max_\|** | **SD(\|E_max_\|)** |
| --- | --- | --- |
| 5-chan planar | 139.60 V/m … 594.91 V/m | 76.2031 V/m |
| 6-chan spherical | 132.54 V/m … 475.65 V/m | 64.0348 V/m |
| 12-chan spherical | 223.66 V/m … 512.61 V/m | 50.0212 V/m |

*Note.* Each mTMS coil array was positioned once to optimally stimulate the center of the primary motor ROI for each subject, and, subsequently, stimulation focality was optimized for each of the ROI elements. |E_max_| is the element-wise maximum stimulation strength in V/m that could be achieved with 100 A/µs current rate constraints.

##### 5-chan planar

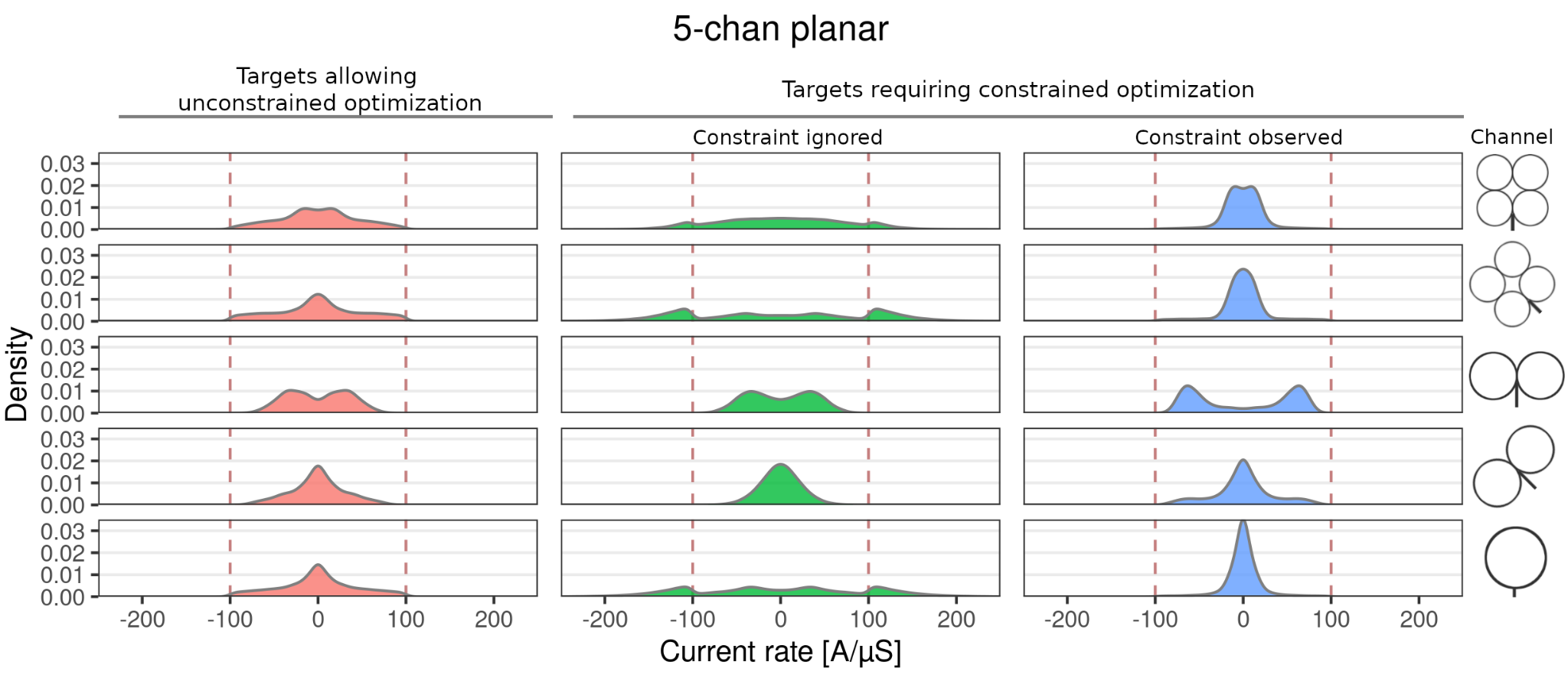

**Figure S9: Constrained and unconstrained optimization: 5-chan planar.** The optimized, single-channel current rates are shown for focality optimized stimulation of each target in the motor cortices of 10 subjects (~12500 targets per subject; one coil array placement) for 100 V/m target strength and 100 A/µS current rate constraint (dashed vertical lines). Left: Subsample of cortical targets that did not require the current constraint to yield focal stimulation at 100 V/m. Center and right columns: Subsample of targets that did require the constraint optimization. For these, results are shown for the case that the current-constraints are ignored (green) or observed (blue).

##### 6-chan spherical

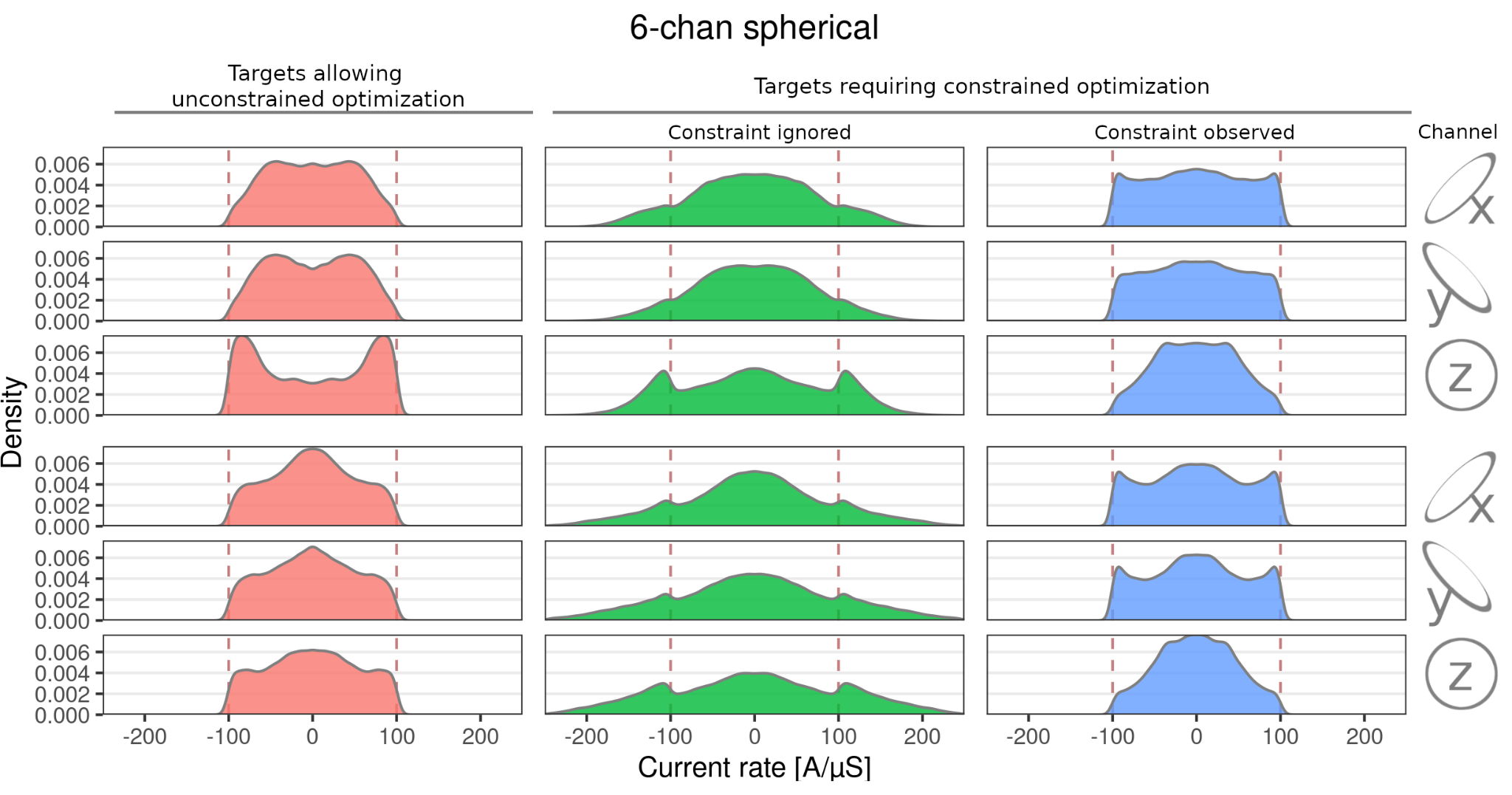

**Figure S10: Constrained and unconstrained optimization: 6-chan spherical.** The optimized, single-channel current rates are shown for focality optimized stimulation of each target in the motor cortices of 10 subjects (~12500 targets per subject; one coil array placement) for 100 V/m target strength and 100 A/µS current rate constraint (dashed vertical lines). Left: Subsample of cortical targets that did not require the current constraint to yield focal stimulation at 100 V/m. Center and right columns: Subsample of targets that did require the constraint optimization. For these, results are shown for the case that the current-constraints are ignored (green) or observed (blue).

##### 12-chan spherical

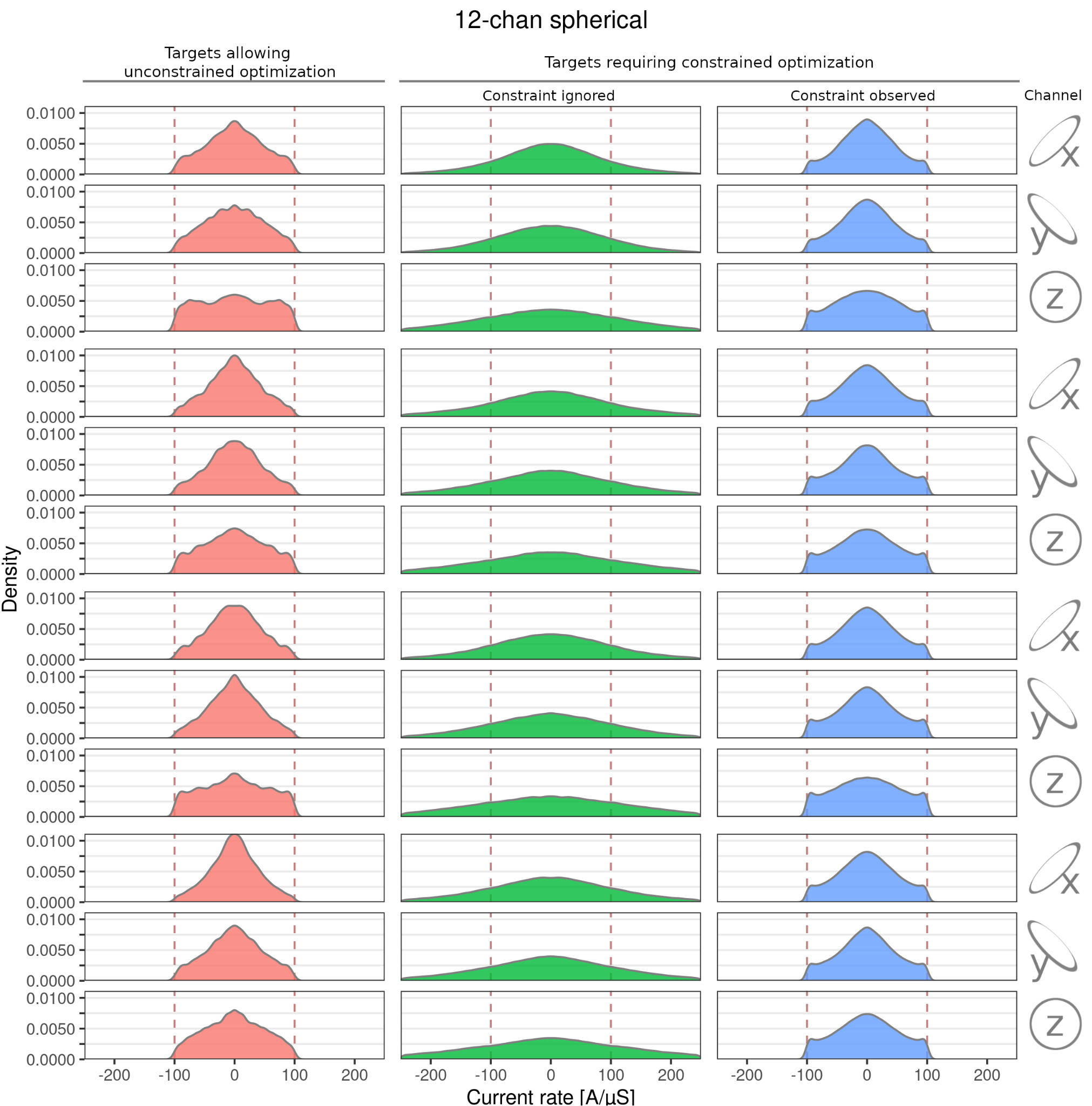

**Figure S11: Constrained and unconstrained optimization: 12-chan spherical.** The optimized, single-channel current rates are shown for focality optimized stimulation of each target in the motor cortices of 10 subjects (~12500 targets per subject; one coil array placement) for 100 V/m target strength and 100 A/µS current rate constraint (dashed vertical lines). Left: Subsample of cortical targets that did not require the current constraint to yield focal stimulation at 100 V/m. Center and right columns: Subsample of targets that did require the constraint optimization. For these, results are shown for the case that the current-constraints are ignored (green) or observed (blue).
